# Surpassing thermodynamic, kinetic, and stability barriers to isomerization catalysis for tagatose biosynthesis

**DOI:** 10.1101/547166

**Authors:** Josef R. Bober, Nikhil U. Nair

## Abstract

There are many enzymes that are relevant for making rare and valuable chemicals that while active, are severely limited by thermodynamic, kinetic, or stability issues (e.g. isomerases, lyases, transglycosidases, etc.). In this work, we study an enzymatic reaction system − *Lactobacillus sakei* L-arabinose isomerase (LsLAI) for D-galactose to D-tagatose isomerization – that is limited by all three reaction parameters. The enzyme has a low catalytic efficiency for the non-natural substrate galactose, has low thermal stability at temperatures > 40 °C, and equilibrium conversion < 50%. After exploring several strategies to overcome these limitations, we finally show that encapsulating the enzyme in a gram-positive bacterium (*Lactobacillus plantarum*) that is chemically permeabilized can enable reactions at high rates, high conversion, and high temperatures. The modified whole cell system stabilizes the enzyme, differentially partitions substrate and product across the membrane to shift the equilibrium toward product formation and enables rapid transport of substrate and product for fast kinetics. In a batch process, this system enables approximately 50 % conversion in 4 h starting with 300 mM galactose (an average productivity of 37 mM/h), and 85 % conversion in 48 h, which are the highest reported for food-safe mesophilic tagatose synthesis. We suggest that such an approach may be invaluable for other enzymatic processes that are similarly kinetically-, thermodynamically-, and/or stability-limited.

---

D-Tagatose is a rare ketohexose sugar with sweetness similar to that of sucrose. However, its glycemic index and caloric value is much lower because of low bioavailability, making it an attractive generally regarded as safe (GRAS) sugar substitute. Recent studies have also demonstrated that it is anti-hyperglycemic ^1^ and prebiotic, which promotes gut health ^2, 3^. Thus, there exists a high demand in food industry for the economical production of rare sugars, like tagatose, exemplified by the 2016 global artificial sweetener market estimated to be USD 3.2 billion. This market is expected to expand given the global diabetes crisis and increasing relevance of prebiotics.

The enzyme L-arabinose isomerase (LAI) that responsible for the reversible isomerization of the pentose L-arabinose to L-ribulose can also isomerize the hexose D-galactose to D-tagatose ^4, 5^. LAI has thus been the enzyme of choice to produce tagatose, although, to date, few commercial bioprocesses exist. A variety of LAIs from different microorganisms have been isolated and have reported optimal activity at a range of temperatures and pH ^4, 6, 7^. Some of the limitations of tagatose biosynthesis using LAI that may be hindering commercial viability are, 1) unfavorable enzymatic kinetics since galactose is not the native substrate of LAI, 2) low enzyme stability, particularly in the absence of divalent metal ions, and 3) low equilibrium constant for galactose to tagatose isomerization.

Few previous reports have been successful at engineering enzymatic properties of LAI for industrial application; often addressing only one of the bottlenecks to productivity. To address the kinetic issue, several groups have used enzyme engineering methods to enhance catalytic efficiency of LAI toward galactose and have shown moderate increases in productivity ^8–13^. To counter low-stability issues, many groups have tested the utility of thermophilic enzymes ^12, 14–16^. However, most thermophilic enzymes rely on divalent metal ions (Mn^2+^, Co^2+^, Fe^2+^) for stability^17^, and high reaction temperatures (≥ 80 °C) result in significant caramelization ^18^, which are all undesirable and must be removed from product, adding to processing costs. Surface-display ^19^ or encapsulation in particles or whole-cells ^20–23^ can stabilize enzymes ^24^. Finally, the thermodynamic limitations of isomerization of galactose to tagatose are severe and, arguably, the most recalcitrant issue since ΔG°_rxn_ ≈ +1.19 kcal/mol ^23^, which indicates theoretical maximum equilibrium conversion ∼ 14% at room temperature. Several approaches have been used to try and overcome this limitation, including the use of an oxidoreductive pathway rather than isomerization ^25^. However, this method resulted in low productivity, byproduct formation, and the need for a sacrificial substrate to regenerate cofactors and drive the reaction uphill. Thermophilic LAI enzymes can achieve higher conversions than their mesophilic counterparts since the equilibrium shifts toward tagatose at higher temperatures ^6^. Whole-cell biocatalysts with GRAS organisms (e.g. lactic acid bacteria (LAB) and *E. coli*) that disproportionately partition substrate and product across their membrane has also been shown to partially circumvent this thermodynamic limitation while simultaneously enhancing enzyme stability; albeit at a kinetic penalty imposed by substrate transport limitations ^26–28^. Recently, cell permeabilization ^28^ and sugar transport overexpression ^29, 30^ were demonstrated as methods to overcome the kinetic penalty imposed by cellular encapsulation.

There have currently been no studies that look to systematically analyze all three limitations – kinetic, thermodynamic, and enzyme stability – of the enzymatic isomerization of galactose to tagatose. This work clearly demonstrates the presence of these three limitations and provides a novel approach to balance their advantages and limitations. We used the food-safe engineered probiotic bacterium *Lactobacillus plantarum* as the expression host due to its increasing relevance to biochemical and biomedical research ^31, 32^. This approach enabled ∼ 50 % conversion of galactose to tagatose in 4 h (productivity of ∼ 38 mmol tagatose L^-1^ h^-1^) ultimately reaching ∼ 85% conversion after 48 h at high galactose loading (300 mM) in batch culture. This is among the highest conversions and productivities reported to date for tagatose production using a mesophilic enzyme. Such an approach is expected to be applicable to other biocatalytic systems where similar trade-offs between kinetics, thermodynamics, and/or stability pose hurdles to process development.

## Results

### Production of tagatose using *L. sakei* LAI (LsLAI) is limited by poor conversion and low thermal stability

At high substrate loading (400 mM galactose) and 37 °C, the reported optimal temperature for this enzyme ^33^, LsLAI purified from *L. plantarum* exhibited an initial forward reaction (turnover) rate (k_initial_) of 9.3 ± 0.3 s^-1^, which is lower than the maximum reaction rate possible by this enzyme of 17.0 ± 1.3 s^-1^ min^-1^ with its preferred substrate, arabinose, at 400 mM. Increasing the reaction temperature to 50 °C increased the initial reaction rate to 11.2 s^-1^ (Fig. 1a) but was accompanied by rapid enzyme inactivation, which is consistent with previous reports of thermal instability of this enzyme ^33^. The enzyme exhibited first-order degradation with a half-life (t_½_) of approximately 18.5 h at 37 °C (Fig. 1b) and 1.5 h at 50 °C. Indeed, the enzyme retains only ∼ 6 % activity after 6 h at this elevated temperature.

**Figure 1:**
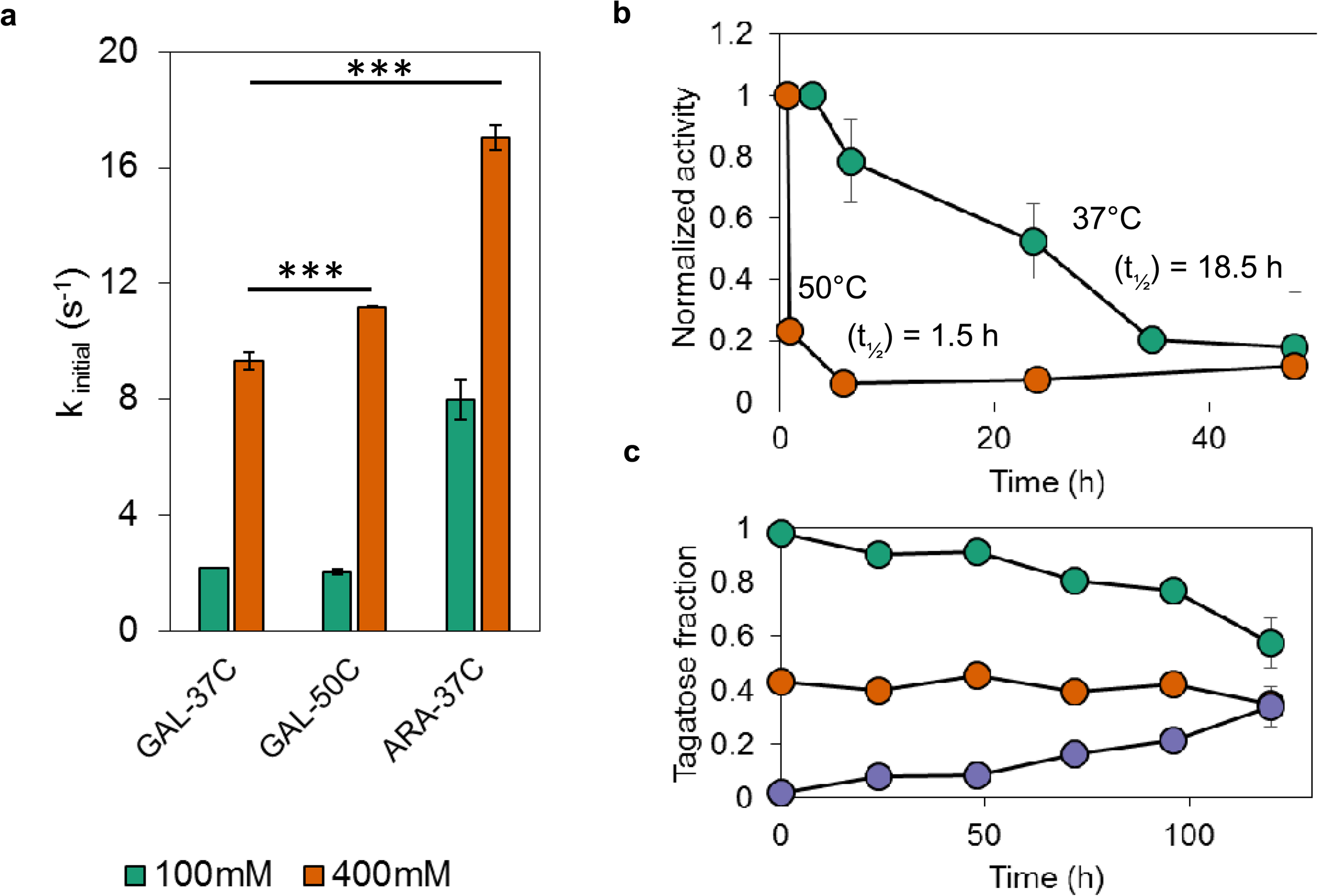
Free-enzyme LsLAI suffers from poor thermal stability, low thermodynamic equilibrium, and kinetic limitations. a) Initial turnover rates of purified LsLAI compared at medium (100 mM, green) and high (400 mM, orange) substrate loading at low (37 °C) and high (50 °C) temperatures. Comparison of the non-native substrate, galactose at low (GAL-37C) and high (GAL-50C) temperature, to that of native substrate, arabinose at low temperature (ARA-37C). b) Loss of activity of purified LsLAI over time incubated at 37 °C (green) or 50 °C (orange). Half-life calculated from first-order decay equation. c) Equilibrium conversion of purified LsLAI starting from 10 mM total substrate with galactose to tagatose ratio at 1:0 (10 mM galactose, purple), 1:1 (5 mM each galactose and tagatose, orange), or 0:1 (10 mM tagatose, green) incubated at 37 °C. Additional enzyme was added every 24 h to account for thermal inactivation. The data are means from three biological replicates. *** = p < 0.001 (Students t-test).

Additionally, the highly reversible isomerization reaction suffers from thermodynamic limitations with reported free-enzyme equilibrium conversions < 50 % (ΔG°_rxn_ = +1.19 kcal mol^-1^) ^23^. Our results indicate similar low conversion of 35 – 40 % after 5 days with pure enzyme at 37 °C with daily supplementation of fresh enzyme to account for inactivation (Fig. 1c).

### *L. plantarum* cell surface display is unsuitable to stabilize LsLAI

To achieve surface localization, we fused LsLAI with six *L. plantarum* surface proteins, which have all been previously described as suitable display carriers, at either the C- or N-terminus (Table S4). The constructs also contained the strong Lp_3050 secretion signal, a 10 amino acid linker containing a thrombin protease cleavage site, and either a C- or N-terminal His_6_ tag for immunofluorescence assays. We characterized surface localization using flow cytometry and measured the whole-cell isomerization activity. All three of the tested C-terminal anchor strains (A1, A2, and A3) exhibited > 90 % surface detection (Fig. 2a) (see Table S3 for strain descriptions). However, the measured tagatose production after a 2 h incubation with 200 mM galactose was only 225 ± 72, 55 ± 39, 91 ± 25 µM tagatose (normalized to individual reaction cell densities of ∼ OD_600_ = 0.5), respectively, significantly less than that of intracellularly expressed (IC1) LsLAI (538 ± 34 µM) (Fig. 2b). LsLAI was detected on N-terminal anchor strains A5 and A6 in 28 % and 96 % of the population and produced 286 ± 31 and 422 ± 34 µM tagatose respectively. Interestingly, strain A4 exhibited low surface detection, comparable to control (WT & IC1) cells while producing the highest titers of tagatose (533 ± 58 µM) among all the surface display constructs. These data indicate there is not a significant correlation (p > 0.05) between whole-cell activity with the presence of displayed LsLAI on the *L. plantarum* cell surface (Fig. 2c).

**Figure 2:**
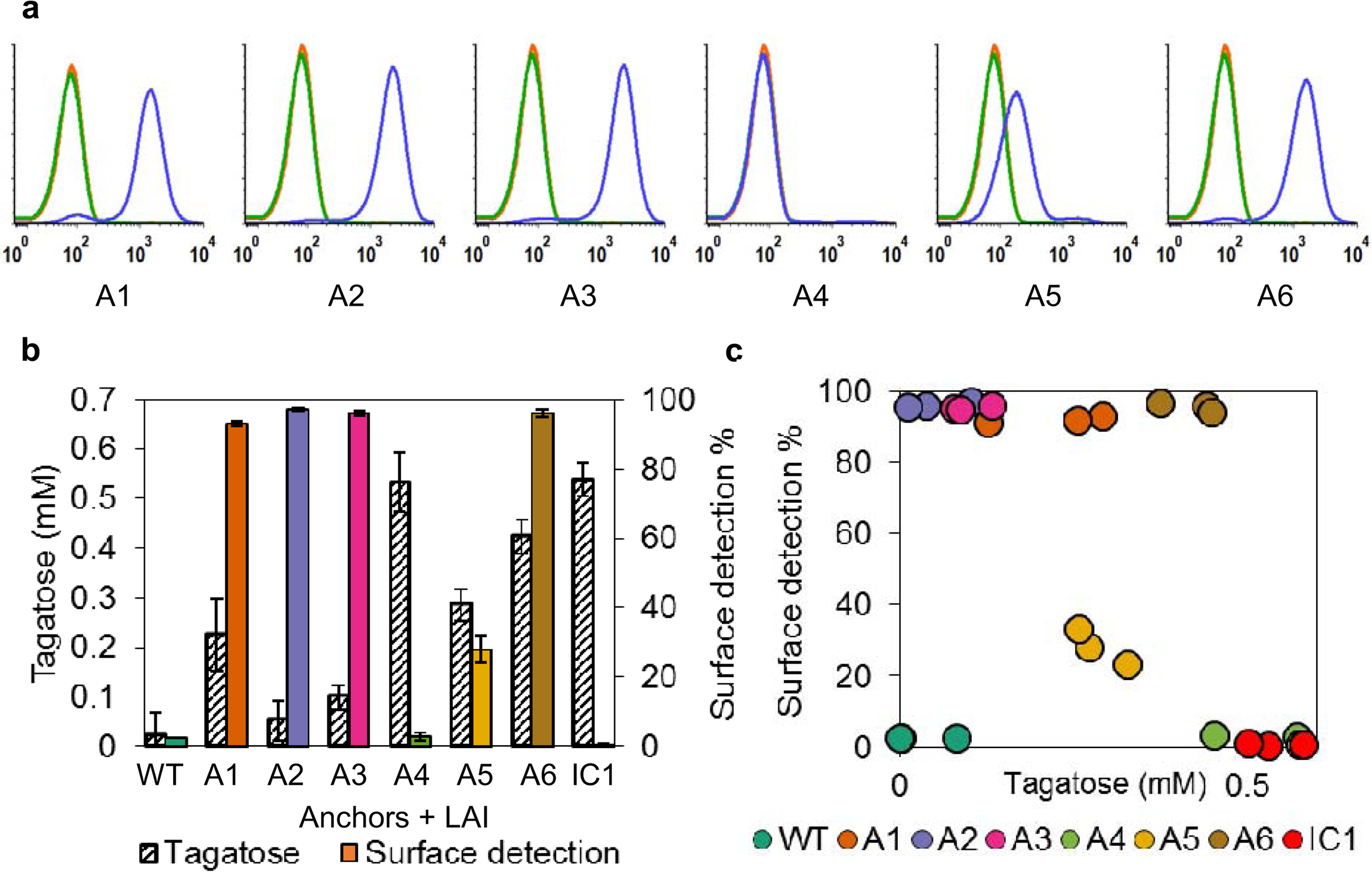
LsLAI surface display and activity. a) Flow cytometry analysis of *L. plantarum* wild-type cells (green), cells expressing intracellular His_6_-tagged LsLAI (orange), or cells expressing LsLAI fused to native surface anchor proteins A1-A6 for display (blue) with 10^5^ counts per sample. Data is normalized to number of counts. b) Comparing the amount of tagatose produced from 200 mM galactose in 2 h (hashed) with the positive percentage of the population with surface detection intensity above that of wild-type “WT” for intracellularly expressed “IC1” or anchor protein A1-A6 (colored) based on negative gate. c) Scatterplot of each replicate measurement correlating surface detection vs. tagatose produced. There is no significant correlation between the display level and activity based on Pearson Product Moment Correlation test (Correlation coefficient = −0.395, p = 0.0564). The data are means from three biological replicates.

Next, we sought to explain the observation regarding absence of positive correlation between display level and activity using strain A6. Sodium dodecyl sulfate (SDS) is a strong anionic surfactant that is commonly used to lyse cells through membrane disruption and denature proteins by destabilizing non-covalent bonds. We posited that SDS could be used to extract surface proteins from Gram-positive bacteria without lysing the cells. We incubated strains IC1 and A6 with either PBS (control) or PBS containing 0.5 % SDS, to remove surface displayed LsLAI, and then quantified surface detection and whole-cell activity. As expected, we detected LsLAI only on the surface of PBS-treated control A6 but not the SDS-treated cells by flow cytometry (Fig. S1a). Interestingly, whole-cell activity of both IC1 and A6 increased approximately 3- and 5-fold, respectively, post-SDS-treatment. These data suggest LsLAI is present in both the intracellular compartment and membrane bound of strain A6 with the vast majority of measured whole-cell activity coming from the intracellular fraction. We confirmed the presence of soluble anchor-LAI fusion (Lp_3014-LsLAI) in the intracellular fraction of A6 via Western blot analysis (Fig. S1b).

Additionally, we noticed enhanced whole-cell activity that we attribute to permeabilization/fluidization of the membrane/cell wall due to removal of surface proteins. While this study only focused one of our six explored anchor protein constructs, we suspect that these results are consistent amongst the other LsLAI surface display constructs. Thus, the measured whole-cell activity differing between constructs was likely dependent on the activity of soluble anchor-LsLAI present in the cytoplasm and not due to surface displayed enzyme.

Finally, to further elucidate the discrepancy between activity of intracellular and surface displayed LsLAI constructs we tested the enzymes activity of secreted, unanchored enzyme. Post-induction, we separated whole cells from culture media and compared the activity of each fraction. We observed a 6-fold higher in activity in the whole-cell fraction of the secretion strain compared to wild-type control cells; however, there was no detectable activity in the supernatant of either strain (Fig. S2a). We confirmed the presence of LsLAI in the supernatant and soluble cell fraction via Western blot analysis (Fig. S2b). These data, taken together with our previous experiments, suggest LsLAI is largely in an inactive state after secretion and that any activity is due to accumulation of active enzyme in the *L. plantarum* cytoplasm.

### Cellular encapsulation improves LAI stability and conversion

Since surface display failed as a stabilization mechanism, yielding minimally active enzyme, we evaluated whether whole-cell *L. plantarum* cells encapsulating LsLAI via cytoplasmic expression (referred to as ‘encapsulated LAI’) could be used as a stabilization method instead. Our data show that such encapsulation within *L. plantarum* protected the enzyme from thermal deactivation at 50 °C; the enzyme retained ∼ 85 % of its initial activity after 24 h (Fig. 3a). Interestingly, resting whole cells at 37 °C displayed no loss of activity after 24 h; in fact, we observed a slight enhancement in activity. This observation could be due to changes to cellular morphology or physiology. Cell-encapsulated LsLAI quickly reached an equilibrium conversion of approximately 60 % in 24 h, overcoming the thermodynamic limitation of < 50 % conversion presented in the pure enzyme system (Fig. 3b). We believe this happens because the substrate (galactose) and product (tagatose) differentially partition across the cell membrane. Although initial reaction rates of encapsulated LsLAI at 37 °C and 50 °C with either galactose or tagatose were lower than that of their free-enzyme counterparts, encapsulated LsLAI favored reaction in the forward direction more than in the reverse direction, as determined by initial rates (Fig. S3). At 50 °C the ratio of forward to reverse initial reaction rate for encapsulated LsLAI was 1.8 whereas for the pure enzyme, the ratio was more unfavorable at 0.7.

**Figure 3:**
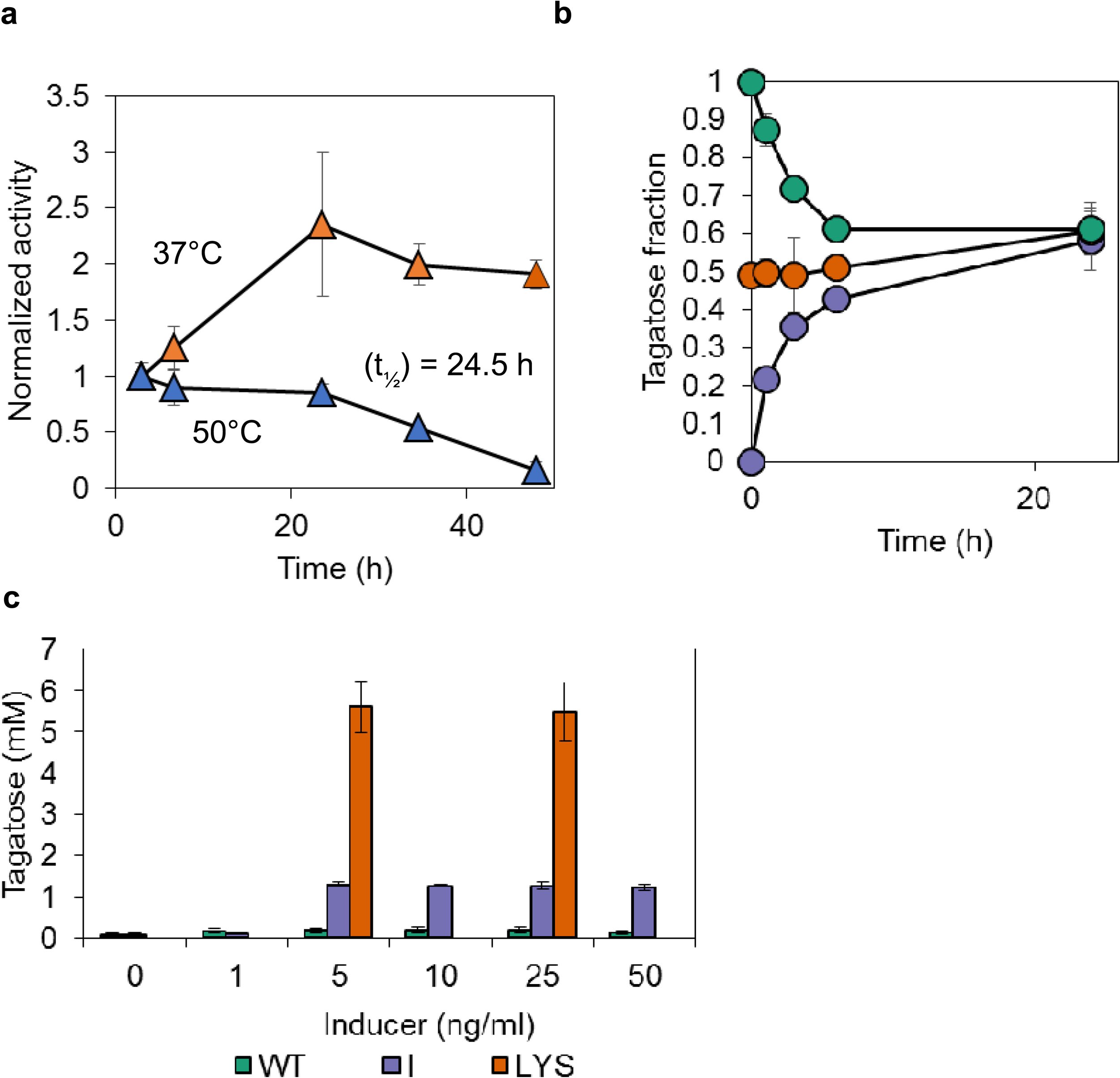
Encapsulation of LsLAI improves equilibrium conversion and provides thermal stability, albeit at a kinetic penalty. a) Loss of activity of encapsulated LsLAI (IC2) over time incubated in PBSM pH 7.4 at 37°C (orange) or 50°C (blue). Half-life calculated assuming first-order decay. b) Equilibrium conversion of encapsulated LsLAI starting from 30 mM total substrate: 1:0 (30 mM galactose, purple), 1:1 (15 mM each, orange), or 0:1 (30 mM tagatose, green) galactose:tagatose respectively. c) Tagatose production from 200 mM galactose of *L. plantarum* wild type “WT” cells (green), IC2 cells expressing LsLAI intracellularly “I” (purple), or crude lysate of the same IC2 cells expressing LsLAI cytoplasmically “LYS” (orange) at different induction concentrations. The data are means from three biological replicates.

To further demonstrate that the *L. plantarum* cell acts as a selective barrier, we tested the capacity of other sugars to inhibit the rate controlling step of galactose to tagatose isomerization. Tagatose production by cell encapsulated LAI was significantly inhibited in the presence of glucose while the presence of arabinose had no effect (Fig. S4). Conversely, glucose had minimal inhibition on tagatose production in pure enzyme reactions while arabinose (the preferred substrate for LAI) significantly inhibited the production of tagatose. These data suggest that the rate-controlling step in whole cell system and pure enzyme are completely different – sugar transporters in whole cell systems and passive diffusion in pure enzyme systems. Further, since glucose is a highly preferred substrate of *L. plantarum*, the fact that glucose inhibits whole cell galactose isomerization suggests that the rate-controlling step is one or more glucose-preferring hexose transporter(s).

While encapsulation was shown to provide thermal stability and a thermodynamic advantage, it imposes a kinetic penalty due to transport limitations across the cell membrane. To test this, we measured initial whole-cell activity under a range of induction levels to determine the maximum achievable activity and compared the whole-cell activity to that of the cell lysate at the same conditions. Strain “I” produced the maximum amount of tagatose (1.5 mM) in 2 h at inducer concentrations ≥ 5 ng/mL (Fig. 3c), indicating that this represented that the system is at maximum reaction rate under all these inducer levels. Conversely, lysate of the same cells induced at 5 and 25 ng/mL inducer conditions generated approximately 4-fold greater tagatose than that of the equivalent whole-cell catalysts in the same 2 h interval. This further supports the idea that cellular encapsulation changes the rate-limiting step to transport and imposes a ceiling on reaction kinetics.

### SDS permeabilized cells overcome all three barriers to tagatose production

Seeking to overcome the transport-limited whole-cell production of tagatose, we investigated the use of permeabilization surfactants − Triton X-100 and SDS. Following the same optimized protocol as previously reported for permeabilization of *L. plantarum* with 1 % Triton X-100 ^28^, we observed a 4.2 ± 0.1 fold increase in tagatose production as compared to untreated cells under the tested conditions (Fig. S5a). We assessed the ability of 0.005 % – 0.5 % SDS to permeabilize cell encapsulated LsLAI and enhance tagatose production. Our results showed the optimal concentration of SDS that enhanced tagatose production via cell permeabilization while still retaining LsLAI intracellularly to be 0.01 % (Fig. S5b). Tagatose production was enhanced 5.7 ± 0.1 fold after permeabilization with 0.01 % SDS, greater than that of treatment with 1% Triton X-100 under the tested conditions. Additionally, this concentration of SDS is below the critical micellular concentration (CMC = 0.24 – 0.26 % at 35 – 40 °C) therefore producing minimal frothing as compared to 1 % Triton X-100, the CMC of which is reported to be 0.05 % ^34^. Although SDS is commonly used to denature proteins, low concentrations of SDS (0.001 – 0.005 %) has been previously used to permeabilize *E. coli* – albeit less successfully than with CTAB (cetyl trimethylammonium bromide) and Triton X-100 ^35^.

To further understand the effect of SDS-treatment on cell permeability, we measured the fluorescence generation rate using 5-carboxyfluorescein diacetate (cFDA) as a substrate. cFDA is an uncharged ester that becomes fluorescent upon hydrolysis by abundant and nonspecific intracellular esterases. Hydrolysis also results in entrapment of the fluorescent dye in the cytoplasm, unless the membrane has been damaged. The rate of generated fluorescence serves as a proxy to determine substrate transport rates assuming hydrolysis rates are higher than transport rate. Wild-type *L. plantarum* treated with 0.005 %, 0.01 %, and 0.05 % SDS had 1.6-, 3.0-, and 3.4-fold increase in substrate transport rates, respectively, compared to untreated cells (Fig. S6). SDS concentrations > 0.5 % resulted in fluorescence leakage, a sign of damaged cell membrane that may also be expelling other cytoplasmic content. These data suggest that the treatment with 0.01 % SDS maintains cellular integrity while increasing tagatose synthesis rate due to a decreased kinetic penalty presented upon encapsulation.

To test whether the enhanced transport could be attributed to changes to the Gram-positive cell wall composition, we mildly treated *L. plantarum* encapsulated LAI (strain IC2) with lysozyme, an enzyme which catalyzes the destruction of the peptidoglycan component of the gram-positive cell wall. These cells mildly displayed a 4.3-fold increase in activity like cells chemically treated with SDS detergent compared to untreated cells (Fig. S7 and Fig. S5). These data suggest that the cell wall poses a transport barrier and SDS treatment may be modifying the structure/rigidity of the cell wall to allow faster transport; albeit further direct experimentation should be done to confirm this hypothesis. However, transmission electron microscopy (TEM) of SDS-treated *L. plantarum* cells found no evidence of pore formation or leakage of cytoplasmic biological components (Fig. S9).

Finally, we monitored the tagatose production with different LsLAI catalyst preparations at high loading (Strain IC2 OD_600_ = 40 or equivalent 0.6 mg/mL purified LsLAI) in batch processes starting with 300 mM galactose. Consistent with our initial experiments, the pure enzyme preparation was limited to 39 ± 2.3 % conversion after 48 h at 37 °C (Fig. 4). Increasing the temperature of the reaction to 50 °C allowed for high initial production of tagatose; however, the enzyme denatured quickly reaching a maximum conversion of only 16 ± 2.8 %. Bacterial encapsulation of LsLAI (IC2 + PBS) produced 47 ± 3.3 % tagatose at 37 °C, improvement over the pure-enzyme system. Encapsulation also provided thermal stability to the enzyme allowing for sustained activity at 50 °C increasing both the rate of production (46 ± 2.3 % in 6 h) and equilibrium conversion (83 ± 6.1 %) compared to unmodified whole-cell catalyst at 37 °C. Rate of production was further increased by modifying the whole-cell catalyst through SDS permeabilization (IC2+SDS) to achieve 59 ± 3.5 % conversion in 6 h, ultimately reaching a similar equilibrium conversion of 85 ± 6.7 % as unpermeabilized cells. Thus, permeabilized, cell-encapsulated LAI overcomes the three reaction limitations by supporting greater thermal stability, higher equilibrium conversion, and faster reaction rates than possible by a free enzyme system.

**Figure 4:**
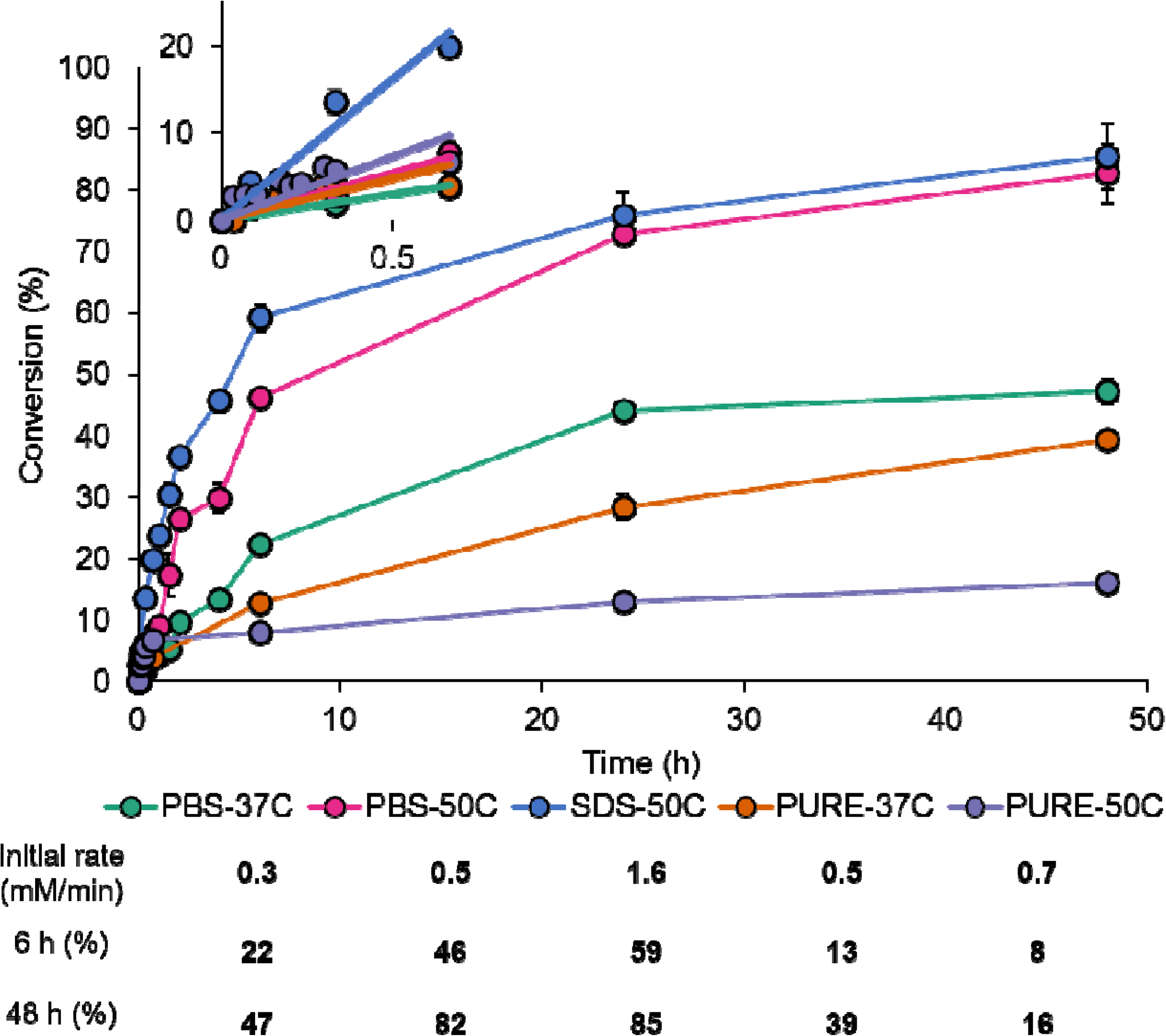
Modified encapsulation of LsLAI maximizes tagatose production. Batch tagatose production over time starting from 300 mM galactose. Encapsulated LsLAI (IC2) was tested for tagatose production as unmodified (PBS) or SDS permeabilized (SDS) compared to purified enzyme at 37 °C or 50 °C to demonstrate its advantages. Encapsulated LsLAI is “PBS-37C” (green), encapsulated LsLAI at 50 °C is “PBS-50C” (pink), permeabilized encapsulated LsLAI at 50 °C is “SDS-50C” (blue), purified free-enzyme LsLAI at 37 °C is “PURE-37C” (orange), and purified free-enzyme LsLAI at 50 °C is “PURE-50C” (purple). Inset plot shows 0 - 1 h data to highlight initial reaction rates of each catalyst for comparison purposes. Salient features are summarized in table. The data are means from three biological replicates.

To investigate the potential to increase the average productivity of our system, we increased the substrate level from 300 mM to 600 mM and 1 M galactose and measured the production of tagatose using permeabilized *L. plantarum* encapsulating *L. sakei* LAI (strain IC2 + SDS) over the course of 24 h. Higher substrate loading lowered the final conversion of the system to 47 % and 39 % at 600 mM and 1 M galactose respectively (Fig. S8). We observed the highest productivity at low conversions, as expected. After 6 hours, the highest average productivity of our system was 9.3 g/L/h of tagatose starting from 1 M galactose substrate loading compared to 5.3 g/L/h starting from 300 mM galactose. Based on these results we conclude that increasing the substrate loading increases average productivity at the cost of conversion, which is consistent with many previous studies ^19,36,37^.

## Discussion

While an increasing number of enzymes are being used for biocatalysis, poor stability under reaction conditions, and unfavorable kinetic properties toward non-native substrates can limit their utility toward biosynthetic functions. Further, for certain classes of enzymes like isomerases, transglycosidases, and lyases, limitations of endergonic or mild exergonic thermodynamics are additional hurdles to economical processes. D-Tagatose synthesis from D-galactose using the enzyme LAI (L-arabinose isomerase) suffers from all three of these limitations. Our results confirm the presence of all three limitations (stability, thermodynamic, kinetic) during enzymatic conversion of galactose to tagatose using the acid-tolerant mesophilic LAI from *Lactobacillus sakei* (LsLAI). At low temperatures (37 °C), the enzyme is stable and quickly reaches equilibrium. However, the product titer is low due to low equilibrium conversion (36 − 39 %) at low temperatures (this study and previously reported ^33^). At elevated temperatures (50 °C), the initial reaction rates are faster, but the enzyme quickly inactivates and is unable to realize the higher conversion and product titer possible at this temperature. These limitations of LAI for tagatose synthesis have been realized previously, and each issue has been addressed individually. While some successes have been reported, especially by selecting or engineering enzymes with more desirable properties ^4, 11, 38–41^, this is the first study to address and overcome all three issues simultaneously.

LAI display on *Bacillus subtilis* endospore surface was shown to improve thermostability while maintaining free enzyme-like kinetic properties ^19^. Therefore, this approach addresses the stability limitations but not the thermodynamic issue. To assess whether cell surface display on *L. plantarum* could improve stability of LsLAI, we tested six previously reported anchor proteins and found there was no correlation linking whole-cell activity and surface display level/accessibility. We showed evidence for presence of cytoplasmic LsLAI-anchor fusions and concluded that enzymatic activity seen was largely due to this active fraction and that the enzyme displayed on the surface was largely inactive. When we tested secretion constructs (i.e., LsLAI with secretion signal but no anchor), we found similar results, suggesting that LsLAI cannot fold extracellularly upon secretion through this pathway and that active enzyme accumulated cytoplasmically. This is surprising since there is no precedence for such an observation in literature. *L. plantarum* utilizes the Sec-pathway to secrete proteins ^42^ and it is generally accepted that translocation occurs co-translationally^43, 44^; therefore, for a protein to be active it must fold properly in the extracellular environment. While we detected LsLAI on the surface using flow cytometry and Western blot analysis, we could not correlate display level to activity or subsequently attribute any detectable activity to the displayed protein at all. Our results suggest the need to be cautious when attributing activity of surface displayed proteins, even if they are readily detected. There are other reports of surface displayed enzymes or antigens have shown lack of correlation between surface detection and activity both in-vitro and in-vivo ^45–47^. We suggest future studies to more directly attribute activity to displayed proteins and not due to presence of active cytoplasmic proteins in whole or lysed cells. We posit that overexpression of surface directed proteins may yield a “back-up” resulting in translation of active protein fusions directly in the cytoplasm. Surface display of LsLAI has been shown before with *B*. *subtilis* endospores; however, the construct is assembled on the spore cytoplasmically ^19^, without the need for any secretion. Our results contradict reports from another study that claimed secretion of active LsLAI using *Lactococcus lactis* ^1^. However, closer analysis of the study revealed that measured activity was never attributed directly to culture supernatant. Rather activity was detected either for whole cells or in the elution fraction post-chromatographic purification, which could have allowed for refolding to occur. Thus, we posit that cell-surface display is not a viable option to improve the stability of this enzyme.

Cellular or abiotic encapsulation has shown to improve enzyme stability against thermal, pH, solvent, and impurity interactions ^24, 48^. In the context of tagatose production, whole cells expressing L-AI has also shown to improve conversion by preferentially partitioning substrate and product across the *E. coli* membrane barrier through active transport ^26^; however, the same was not observed in resting *L. lactis* cells ^49^. Our results demonstrate that encapsulation of LAI in *L. plantarum* behaves in a manner similar to that in Gram-negative *E. coli* but not Gram-positive *L. lactis*. That is, the cell membrane preferentially allows passage of galactose to the intracellular space relative to tagatose to aid the forward reaction. A concurrent advantage of the encapsulation was that the enzyme stability was markedly improved. Specifically, while the free enzyme rapidly inactivates at 50 °C, the cell-encapsulated enzyme maintains activity for a prolonged period. Indeed, the resistance to lysis at elevated temperatures is afforded by the thick cell wall of the Gram-positive *L. plantarum*, as opposed to Gram-negative *E. coli*, which is known to readily lyse at 50 °C ^50^. Thus, while many previous studies have demonstrated the benefits of using whole cells to enhance enzyme stability, this work is the first to show the twofold benefit by combining improved stability with substrate-product partitioning. While the selective membrane circumvents the thermodynamic barrier that limits maximum conversion at moderate temperatures, it imposes a barrier to mass transport – a penalty to reaction kinetics. We addressed this issue next.

Cell permeabilization using chemical agents like detergents or solvents have been shown to non-selectively increase substrate exchange while maintaining biological cellular components ^51, 52^, although the exact mechanism of this phenomenon is poorly understood. We used detergents to improve reactant/product transport across the *L. plantarum* cell membrane without altering its selectivity. In our best-case scenario, sub-micellar concentration of SDS resulted in significant increase in whole cell reaction rates to free-enzyme-like rates while still allowing the system to circumvent the thermodynamic barrier to product conversion. Since the conversion was unchanged, we presume that the detergents were not making pores in the cell membrane. TEM images of SDS-treated and untreated cells further confirmed that the cell wall and membrane were intact under reaction conditions. Interestingly, permeabilization increased the initial reaction rate in the forward direction 3.8-fold, however the rate of the reverse reaction remained low, indicating that the membrane retained its selectivity (Fig. S3). Studies with cFDA further confirmed that cytoplasmic components were not “leaking” into the extracellular space and that the cellular integrity was maintained. Thus, substrate and product transport were still likely facilitated through transporters, as in untreated cells, which is also consistent with the observation that glucose inhibits only whole cell reactions (but not that of free enzyme) and conversion reaches the same maximum with or without detergent treatment. Since SDS and Triton-X treatment resulted similar outcomes, we believe there are no specific charge-related effects of these detergents, either. Adding SDS to a pure enzyme solution did not alter its kinetic properties, indicating that the detergents do not directly interact with the enzyme in any appreciable manner. However, our study revealed that treatment with SDS removed extracellular- and surface-attached proteins, potentially altering the fluidity of the membrane and/or permeability of the cell wall. The latter is supported by the fact mild lysozyme treatment also increases reaction rates, suggesting that the thick Gram-positive peptidoglycan poses a transport barrier.

Having addressed all three limitations (stability, thermodynamics, kinetics), we found that *L. plantarum* whole-cell biocatalyst achieved a conversion from of ∼ 50 % in 4 h and ∼ 85 % 48 h in batch process at 50 °C starting with 300 mM galactose. Improved stability is apparent by comparison with the free enzyme system, which quickly inactivates at 50 °C and achieves only 16 % conversion in 48 h. Enhanced equilibrium conversion compared to free enzyme system can be seen at both 50 °C and 37 °C, but especially in the latter, where the free enzyme is more stable but is still able to achieve only 13 % conversion in 4 h and 47 % in 48 h. Finally, improved kinetics can be seen compared to unpermeabilized cells where initial reaction rates at 0.5 mM/min compared to 1.6 mM/min in permeabilized cells. This rate is higher than the maximum reaction rates sustainable by the free enzyme for extendable period (initial rate at 37 °C for free enzyme = 0.3 mM/h) due to the ability to operate at 50 °C. This is the among the highest reported average productivity (5.3 g/L/h after 6 h) for biocatalytic tagatose production, and the highest conversion reported for a mesophilic enzyme (∼85 %) ^19, 53^. Average productivity of the modified cell encapsulated LAI batch process could be improved by increasing the initial concentration of galactose from 300 mM to 1 M, albeit with lower conversion. Although further improvements to conversion may be possible by including borate in the reaction buffer, demonstrated previously, it would also increase purification costs ^16, 27, 49^. Further, this system nearly matches the conversion obtained in a recent study with a extremophilic LAI at 95 °C and pH 9.5 that required use of Ni^2+^ and borate ^54^ but at 50 °C and without use of toxic metals and salts. Additionally, the system avoids product caramelization, which is known to happen at high temperatures (≥ 80 °C) ^18^. Thus, the system described here demonstrates how to overcome three major limitations of enzymatic catalysis (stability, thermodynamic, kinetic) for tagatose production using LAI. We believe that a similar strategy may be applicable to other biocatalytic systems as well – particularly lyases ^55^, transglycosidases ^56^, and isomerases ^17, 26, 36, 57^ – where free enzyme performance is poor due to stability, thermodynamic, and/or kinetic limitations.

## Statistics and data reproducibility

All the experiments were conducted using biological replicates and were carried on different days as specified to calculate measure of variability between the samples. All the data shown are mean with error bars representing standard deviation.

## Acknowledgments

This work was partly supported by National Institutes of Health (NIH) Grant DP2HD91798 and R03HD090444 to N.U.N. We thank Dr. Michiel Kleerebezem (Wageningen University & Research, Wageningen, Netherlands) for sharing strain *L. plantarum* WCFS1 and Dr. Lars Axelsson (Nofima, Tromsø, Norway) and Dr. Geir Mathiesen (Norwegian University of Life Sciences, Ås, Norway) for sharing plasmids. Electron microscopy was conducted utilizing the W.M. Keck Foundation Biological Imaging Facility at the Whitehead Institute. Special thanks to Nicki Watson for performing the electron microscopy imaging.

## Author contributions

J.R.B. and N.U.N. planned the experiments, wrote the manuscript, and analyzed the results. J.R.B conducted all the experiments.

## Materials and Methods

### Reagents and Enzymes

All enzymes for cloning were purchased from NEB (Beverly, MA). DNA primers were ordered from Eurofins Genomics LLC (Louisville, KY) or GENEWIZ Inc. (Cambridge, MA). Growth media and chemicals were purchased from Amresco (Solon, OH) or RPI Corp (Mount Prospect, IL). High-purity L-arabinose and D-galactose were purchased from Sigma-Aldrich (St. Louis, MO). Plasmid Mini Kit I, PCR Purification and Gel Extraction Kits were obtained from Omega Bio-tek (Norcross, GA). Mouse-anti-His_6_ primary antibody and goat anti-mouse IgG H&L (Alexa Fluor® 488) were purchased from Thermo Fisher Scientific (Waltham, MA).

### Bacterial Strains and Culture Conditions

*Lactobacillus plantarum* WCFS1 (kind gift from Prof. Michiel Kleerebezem, Wageningen University & Research) was cultivated in deMan−Rogosa−Sharpe (MRS) medium at 30 °C without agitation. *E. coli* strain NEB5α (New England Biolabs, Ipswich, MA, USA) was grown in Luria−Bertani (LB) medium at 37 °C with shaking at 225 rpm. When needed, media were supplemented with erythromycin at concentrations of 5μg/mL and 200 μg/mL for *L. plantarum* and *E. coli*, respectively. Solid media were prepared by adding 1.5 % (w/v) agar to the respective media.

### Plasmid and Strain Construction

DNA amplification was performed with Phusion High-Fidelity DNA Polymerase according to the manufacturer’s instructions. *E. coli* was used as host for plasmid propagation before transformation into *L. plantarum*. The amplified sequences were verified by a commercial sequencing service (GeneWiz). A list of all strains and plasmids used in this study in Table S2 and Table S3.

*L. plantarum* surface display plasmids were constructed based on the previously published pSIP401 based plasmid pLp_3050Ag85B-E6cwa2 (Table S2). The LAI gene from *L. sakei* 23K was synthesized by Twist Biosciences. This gene was subsequently cloned using primer pair oJRB133 and oJRB134 and inserted into pLp_3050Ag85B-E6cwa2 using BfaI and HindIII restriction sites to create plasmid pJRB01Q-LSH. pJRB02Q-LSH was created via was the amplification of Lp_2162 from the *L. plantarum* chromosome using primer pair oJRB10 and oJRB11 and inserted into pJRB01Q-LSH using MluI and HindIII restrictions sites. pJRB03Q-LSH was created via was the amplification of Lp_2940 from the *L. plantarum* chromosome using primer pair oJRB20 and oJRB21 and inserted into pJRB01Q-LSH using MluI and HindIII restrictions sites. pJRB05Q-LSH was created via the amplification of Lp_1261 from the plasmid pLp_1261Ag85B-E6 (Table S2) using primer pair oJRB7 and oJRB8 before being overlapped with *L. sakei* LAI and inserted into pJRB01Q-LSH using SalI and PmlI restriction sites. pJRB06Q-LSH was created via amplification of Lp_1452 from the *L. plantarum* chromosome using primer pair oJRB14 and oJRB15 and inserted into pJRB05Q-LSH using SalI and PmlI restrictions sites. pJRB08Q-LSH was created via the amplification of Lp_3014 from the *L. plantarum* chromosome using primer pair oJRB82 and oJRB83 and inserted into pJRB05Q-LSH using SalI and PmlI restrictions sites. pJRB04Q-LSH was created via amplification of the *L. sakei* LAI gene using primer pair oJRB170 and oJRB171 and inserted into pJRB01Q-LSH using NdeI and HindIII restriction sites. pJRB09Q-LSH was created via amplification of the *L. sakei* LAI gene using primer pair oJRB304 and oJRB305 and inserted into pJRB01Q-LSH using SalI and PmlI restriction sites. Plasmid pJRB14Q-LSH was created via amplification of the *L. sakei* LAI gene and 6x-His tag using primer pair oJRB457 and oJRB458 and inserted into pSIP411 (Table S2) using NcoI and HindIII restriction sites.

Each plasmid was transformed into chemically competent *E. coli* NEB5α cells according to the manufacturer’s instructions. *L. plantarum* was transformed using a previously described electroporation method ^58^ with slight modifications. Briefly, stationary phase cells were subcultured to an OD_600_ = 0.1 in MRS and allowed to grow until an OD_600_ = 0.85. Cells were harvested via centrifugation at 4 °C and washed twice in ice-cold 10 mM MgCl_2_ and once with 0.5 M sucrose and 10 % glycerol. Cells were resuspended in 1:50 (v/v) of the same solution and kept on ice prior to transformation. 100 ng of plasmid DNA was diluted into 5 μL of water and mixed with 50 μL electrocompetent cells immediately before adding to a chilled 1 mM electroporation cuvette. The bacteria were transformed using a BioRad Pulser at 1300 V with a fixed time constant of 5 ms. 1 mL of warm MRS was added and the cells transferred to a new centrifuge tube and allowed to recover statically at 37 °C for 3 h before plating on appropriate media containing selective antibiotic. Positive transformants were confirmed using colony PCR.

### Enzyme Purification

*L. plantarum* cells carrying pJRB14Q-LSH were grown and induced as described previously. Cells were washed in lysis/equilibration buffer (50 mM sodium phosphate, 300 mM sodium chloride, 15 % glycerol, pH 8.0) and resuspended in lysis buffer containing 10 mM lysozyme and incubated at 37 °C with shaking for one hour. Lysis was achieved via sonication and soluble protein collected as described previously.

Purification was achieved via immobilized metal affinity chromatography (IMAC) using an N-terminal His_6_ tag. TALON metal affinity resin (Invitrogen) was used as directed by the manufacturer. A single elution of protein was achieved in elution buffer containing 500 mM imidazole. Active fractions were pooled, and the buffer exchanged with PBS containing 1 mM MgCl_2_ (PBSM) using a Microsep 10 kDa Omega centrifugation spin column (PALL Corp; Port Washington, NY). Purification was confirmed via SD-PAGE analysis. Purified enzyme was stored in PBSM with 20 % glycerol at −80 °C prior to use.

### Enzyme Analysis

*L. plantarum* wild-type (WT) cells or those carrying plasmids pJRB01Q-LSH (A1), pJRB02Q-LSH (A2), pJRB03Q-LSH (A3), pJRB04Q-LSH (IC1), pJRB05Q-LSH (A4), pJRB06Q-LSH (A5), pJRB08Q-LSH (A6) were cultured in MRS media with antibiotic as needed at 37 °C overnight. Overnight cultures were diluted 1:50 (approximate OD_600_ = 0.1) in MRS media with antibiotic as needed and grown at 37 °C for 2.5 h. Cells were then induced with IP-673 (synthesized by Life Technologies; Carlsbad, CA) at a final concentration of 25 ng/ml and MgCl_2_ was added to a final concentration of 1 mM to support enzyme folding. Cultures were incubated at 37 °C for an additional 3 h before harvesting the cells by centrifugation at 3000 × g for 5 min.

Cell pellets were washed in 1 mL of 20 mM phosphate buffer (PB) pH 6.8 and transferred to a 1.7 mL centrifuge tube. Cells were washed twice with 1 mL of PB before being suspended in 200 μL PB buffer. The enzyme activity assay was performed in a 96-well plate as follows: 20 μL cell suspension was incubated with 0.8 mM MnCl_2_, 0.8mM MgCl_2_, sugar (200 mM galactose or 200 mM glucose) and diluted to 200 μL with PB. The cell density within each well was measured for data normalization at the start of the reaction. The plates were sealed and incubated at 37 °C for 2 h. Enzyme activity was measured colorimetrically using the cysteine− carbazole−sulfuric-acid method (CCSAA) ^59^.

Absorbance at 560 nm was measured after 1 h of incubation at room temperature. Triplicate technical replicates of each cell suspension were analyzed and averaged due to variability in the assay itself.

To determine the conversion achieved with our balanced system, *L. plantarum* cells carrying pJRB14Q-LSH (IC2) and wild-type were grown and induced as described previously with slight modification. Cells were induced with 25 ng/mL IP-673 for 8 h instead of 3 h to maximize the cell density of the culture. Cells were harvested via centrifugation and washed twice in phosphate-buffered saline pH 7.4 with 1.0 mM MgCl_2_ (PBSM). Cells to be permeabilized were diluted to an OD_600_ = 3.0 before incubation with 0.0 1% SDS in PBSM for 30 min at room temperature. The enzyme activity assay was performed in 5 mL tubes as follows: cell suspensions adjusted to an OD_600_ = 40 were incubated in phosphate-buffered saline (PBS) pH 7.4 with 1.0 mM MgCl_2_, 300 mM galactose and adjusted to 5 mL with H_2_O. The conversion and productivity at increasing concentrations of galactose was performed by increasing the substrate concentration to 600 mM or 1 M with the same assay conditions.

Initial rates of reaction for both pure-enzyme, unmodified, and modified whole-cell catalysts were determined. *L. plantarum* cells carrying pJRB14Q-LSH were grown and induced for 8 h as described previously. Permeabilized cells were prepared as described previously. The enzyme activity assay was performed in 200 μL PCR tubes as follows: purified enzyme or cell suspensions were incubated in PBSM, variable substrate concentration of galactose or tagatose, and adjusted to 100 μl with H_2_O. Reactions were initiated by adding 1/10^th^ volume cell suspensions adjusted to an OD_600_ = 10 or 2 μg of purified enzyme to prewarmed reaction mixtures in a PCR block at either 37 or 50 °C. Reactions proceeded for 6 min before termination to measure initial reaction rates. Cell suspensions were immediately centrifuged for 1 min using a benchtop quick-spin centrifuge and the supernatant was transferred to a 96-well 0.2 μm centrifugal filter plate with collection plate. The remaining solution was filtered via centrifugation and stored at −80C until analysis. Purified enzyme reactions were terminated by adding 1/10^th^ volume 2M perchloric acid. Prior to HPLC analysis the reactions were neutralized using a sodium hydroxide PBS buffered solution. “k_initial_” is defined as the initial reaction rate of each catalyst at a given, non-saturating substrate concentration and stopped the reaction prior to 10% total conversion.

Enzyme stability was measured by incubating either purified enzyme or unmodified whole cell catalysts in PBSM at 37 or 50 °C statically for the duration of the experiment. At determined time intervals and aliquot of catalyst was added to PBSM reaction mixture containing 300mM galactose at the same temperature as the catalyst. Reactions proceeded for 20 min before termination as described previously. Tagatose production was confirmed via HPLC analysis.

Equilibrium conversion was measured by in incubating either purified enzyme or unmodified whole cell catalysts in PBSM at 37 °C. Purified enzyme was incubated with either 10 mM galactose, 5 mM of galactose and tagatose, or 10 mM tagatose. The reaction was sampled every 24 h and additional enzyme was added after sampling due to incomplete reaction due to enzyme degradation. Whole cell catalyst was incubated with either 30 mM galactose, 15 mM of galactose and tagatose, or 30 mM tagatose.

The selectivity of the *L. plantarum* cell barrier was tested by comparing the inhibition of the transport of galactose compared to that of the pure enzyme. *L. plantarum* strain IC2 carrying pJRB14Q-LSH was grown and induced for 8 h as described previously. *L. plantarum* encapsulating LAI (IC2) or purified LAI (PURE) was added to reaction mixture containing and incubated at 37 °C. Reactions proceeded for 20 min before termination as described previously.

### HPLC Analysis

Agilent Infinity HPLC system (Agilent; Santa Clara, CA) equipped with a Hi-Plex Ca+ 300 x 7.7mm column with a guard column and detected using Agilent Infinity 1260 ELSD detector. The mobile phase was filtered deionized water run at 85 °C with a flow rate of 0.6 mL/min. The ELSD detector’s evaporation temperature was set at 90 °C, the nebulizer temperature set to 50 °C, and nitrogen flow rate at 1.6 SLM (standard liter per minute). For calculation of the reaction product(s) L-arabinose, D-galactose, L-ribulose, and D-tagatose standards were included in the run.

### Flow Cytometry and Immunostaining

*L. plantarum* WT cells or those carrying plasmids pJRB01QLs-pJRB08QLs were grown and induced as described previously. Cells were collected by centrifugation at 5000 ×g for 5 min and washed twice in PBS before being suspended to an OD_600_ = 1.0. An aliquot of 200 μL cells was pelleted and was resuspended in 200 μL PBSA (PBS with 2 % bovine serum albumin). The suspension of cells rested on ice for 30 min before the cells were pelleted and solution removed. The cells were then resuspended in 50 uL of PBSA containing 1 μg mouse-anti-His_6_ primary antibody. The cells were incubated for 1h at room temperature with gentle rocking. Cells were collected and washed thrice with PBSA before being suspended in 50 μL of PBSA containing 0.4 μg goat anti-mouse IgG H&L (Alexa Fluor® 488). The cells were incubated for one fifteen minutes on ice and blocked from light. Finally, cells were washed thrice in PBSA and resuspended in 500ul of PBSA. A 20ul aliquot was diluted 10X in PBSA and transferred to a flat-bottom 96-well plate for flow cytometric analysis. A positive control sample was used to calibrate the fluorescence intensity. A negative control sample was used to gate the level of autofluorescence of the cells and any non-specific secondary antibody binding (Fig. S10). A total of 1 × 10^5^ cell counts per sample plotted on a ‘logical’ or bi-exponential x-axis. Data analyzed using FCS Express 6 (De Novo Software, Glendale, CA). Surface detection level was calculated as the fraction of cells with a measured fluorescence greater than that of the gated negative control.

### Surface Treatment

*L. plantarum* WT cells or those carrying either pJRB04Q-LSH or pJRB08Q-LSH were grown and induced as previously described. Cells were washed in PBS twice before further processing. A concentrated cell solution was incubated in PBS containing either 0 % or 0.5 % SDS for 1 h before being pelleted by centrifugation. The SDS solution was removed and the cell pellet washed twice with PBS. Surface detection was measured via immunofluorescence flow cytometry as described previously. Whole-cell activity was measured via the CCSAA after a 1-hour incubation in 200 mM arabinose and 1 mM MgCl_2_ in 100 mM sodium acetate buffer pH 5.0 at 37 °C.

### Western Blot Analysis

*L. plantarum* WT cells or those carrying either pJRB04Q-LSH or pJRB08Q-LSH were grown and induced as previously described. A concentrated cell solution was incubated with 0.05 % SDS in PBS for 30 min at 37 °C before being pelleted. The SDS buffer solution was removed and stored separately. The cell pellet was then washed twice with PBS. A fresh cell solution of the same concentration was prepared in PBS as a control. Both aliquots of cells were then incubated with 10 mg/mL lysozyme for 1 h at 37 °C before being lysed via sonication. Sonication was performed using a Branson 150 system equipped with a microtip probe sonicating samples on ice at 55 % amplitude in 30 s ON 2 min OFF cycles for a total of 5 min of ON time. The insoluble cell fraction containing membrane components and the insoluble protein fraction was pelleted via centrifugation at 20,000 ×g for 5 min. The soluble fraction was removed, filtered through 0.4 μm sterile filter, and stored separately. An aliquot of each of the cell fractions for the three cell types was added to denaturing loading dye and incubated at 98 °C for 15 min. 10 μL of each sample were added to a 4 % - 12 % Bis-Tris PAGE gel in MOPS buffer and run at 75 V for 15 min followed by 110 V for 1.5 h. After separation the gel was removed from the cassette and washed twice in deionized water. The proteins were next transferred to a PVDF membrane for Western blot analysis. Transfer was visually confirmed through the transfer of the stained protein ladder. The membrane was blocked overnight at room temperature with 5 % skim milk in Tris buffered saline containing 1 % Tween-20 (TBST). The membrane was washed twice with TBST before being incubated with 0.2 µg/mL primary antibody in blocking buffer for 2 h at room temperature. The membrane was washed twice with TBST before being incubated with 2 ng/mL of the horseradish peroxidase (HRP) conjugated secondary antibody at room temperature for 1 h. The membrane was then washed twice with TBST before incubation in the substrate containing solution (Thermo SuperSignal West) for 15 min at room temperature. The chemiluminescent signal was captured electronically via a camera set to a 30 min exposure time.

### Cell Permeabilization

*L. plantarum* WT cells or those carrying pJRB14Q-LSH (IC2) were grown and induced and washed in PBS as previously described. Cells were resuspended to an OD_600_ = 3.0 in 200 μL of PBS. Samples were either incubated with 1% Triton X-100 ^28^, 0.1 % SDS, 0.1 μM chicken egg lysozyme, or no treatment at room temperature for 1 h. The solution was removed, and the cell pellet washed twice with PBS before enzymatic activity of the whole-cells were measured as described previously using 200 mM galactose in PBS pH 7.4 containing 1 mM MgCl_2_. Activity was quantified via the CCSAA and normalized to cell concentration as described previously.

The optimal concentration of SDS to maximize whole-cell activity was determined. Cells were prepared as previously described. Cell lysate from the equivalent cell density was prepared via lysozyme treatment via sonication as described previously. Permeabilized samples were either incubated with varying concentrations of SDS, or no treatment at room temperature for 30 min. The solution was removed, and the cell pellet washed twice with PBS before enzymatic activity of the whole-cells were measured as described previously using 200 mM galactose. Activity was quantified via the CCSAA and normalized to cell concentration as described previously.

Cell permeabilization was also tested using carboxyfluorescein diacetate (cFDA) (Sigma-Aldrich) as substrate. Cell suspensions were diluted to OD_600_ = 0.5 in PBS in a black polystyrene 96-well plate. The reaction was started by adding 10 mM cFDA to the cell suspensions. Transport of cFDA was estimated by measuring the fluorescent signal released when cFDA becomes activated upon intracellular cleavage via nonspecific esterases. The total fluorescence was measured on a plate-reader using an excitation and emission wavelength of 485 nm and 520 nm, respectively, every 15 s for 30 min. Apparent transport rates were calculated from the slope of the linear fit to the data.

### Electron Microscopy

The cells were fixed in 2.5 % gluteraldehyde, 3 % paraformaldehyde with 5 % sucrose in 0.1 M sodium cacodylate buffer (pH 7.4), pelleted, and post fixed in 1 % OsO_4_ in veronal-acetate buffer. The cells were stained *en block* overnight with 0.5 % uranyl acetate in veronal-acetate buffer (pH 6.0), then dehydrated and embedded in Embed-812 resin. Sections were cut on a Leica EM UC7 ultra microtome with a Diatome diamond knife at a thickness setting of 50 nm, stained with 2 % uranyl acetate, and lead citrate. The sections were examined using a FEI Tecnai spirit at 80 KV and photographed with an AMT ccd camera.

## Data availability

Data supporting the findings of this work are available within the paper and its Supplementary Information files. A reporting summary for this article is available as a Supplementary Information file. The source data underlying Figs. 1, 2, 3, 4, and Supplementary Figures are provided as a Source Data file. All other data are available from the corresponding author upon reasonable request.

