## Supplemental Tables and Figures for "Surpassing thermodynamic, kinetic, and stability barriers to isomerization catalysis for tagatose biosynthesis"

**Supplementary Table 1: Primers used in this study**

| **Primer** | **Description** | **Sequence** |
| --- | --- | --- |
| oJRB7 | Lp_1261_F | GTAAgtcgacAATTTCAAAACAGCTGCAAAAGTAA |
| oJRB8 | Lp_1261_R | AATGgcatgcGCCCCCGTTCTTACCGAGACGGTAT |
| oJRB10 | Lp_2162_F | GTACacgcgtGCTGCCAATGCTGCCTCAAT |
| oJRB11 | Lp_2162_R | GTACaagcttCTACATGGTTGTTTTTTGACTCGT |
| oJRB14 | Lp_1452_F | GCCAgtcgacAAGAAATGGCTCATTGCCCTTGCTGGTGTC |
| oJRB15 | Lp_1452_R | CTAAgcatgcTTGAACCGTGACTTTAGGTTCGTAAGACTTCC |
| oJRB20 | Lp_2940_F | GTACacgcgtGCCGAATCTAACACTCGAACCGG |
| oJRB21 | Lp_2940_R | ATGCaagcttTTAATCAGTTGTTTTATGGCGCCGTTG |
| oJRB34 | Lp_1261ovrlp_R | ttcatgatgGCTGCCGCGCGGCACCAGgcatgcGCCCCCGTTCTTACCGAGACGGTAT |
| oJRB35 | Lp_1452ovrlp_R | ttcatgatgGCTGCCGCGCGGCACCAGgcatgcTTGAACCGTGACTTTAGGTTCGTAAG |
| oJRB82 | Lp_3014_F | TTGCCAgtcgacAAAAAACTTGTAAGTACAATCGTAACT |
| oJRB83 | Lp_3014_R | TGCTAAgcatgcAAGGGCCCAAGCAGCCAT |
| oJRB132 | LsLAIovrlp_F | GCATGAgcatgcCTGGTGCCGCGCGGCAGCctcgagTTAAATACAGAAAATTATGAATTT |
| oJRB133 | LsLAIovrlp_R | GTCTACtctagaTTTAATATTGACGTAAGTCAA |
| oJRB170 | LsLAI_IC1_F | GCATTActagATGTTAAATACAGAAAATTATGAATTTTG |
| oJRB171 | LsLAI_IC1_R | GCATTAaagcttTTAGTGATGATGATGATGATGacgcgtTTTAATATTGACGTAAGTCAAATC |
| oJRB304 | LsLAI_SEC_F | TAGGATgtcgacATGGAACAAAAACTTATTTCTGAAGAAGATCTGtctagaATGTTAAATACAGAAAATTATGAATTT |
| oJRB305 | LsLAI_SEC_R | TCATTAcacgtgTTAATATTGACGTAAGTCAAATCA |
| oJRB457 | LsLAI_IC2_F | TAAGATgccatggtaTTAAATACAGAAAATTATGAATTTTG |
| oJRB458 | LsLAI _IC2_R | CTATTAaagcttTCAATGATGATGATGATGATGTCTAGATTTAATATTGACGTAAGTCAA |

**Supplementary Table 2: Plasmids used in this study**

| **Plasmid** | **Description** | **Reference** |
| --- | --- | --- |
| pLp_3050Ag85B-E6cwa2 | pSIP401 based plasmid containing an oncofetal antigen and Lp_2578 anchor for *L. plantarum* surface display. | ^58^ |
| pLp_1261Ag85B-E6 | pSIP401 based plasmid containing invasion and Lp_1261 anchor for *L. plantarum* surface display. | ^59^ |
| pLp_1452Inv | pSIP401 based plasmid containing invasion and Lp_1452 anchor for *L. plantarum* surface display. | ^59^ |
| pSIP411 | Lactobacillus inducible plasmid system with broad host SH71 origin. | ^60^ |
| pJRB01Q-LSH | pSIP401 based plasmid containing Ls-araA, sppQ, Lp_2578, SP_3050, His_6_ tag | This work |
| pJRB02Q-LSH | pSIP401 based plasmid containing Ls-araA, sppQ, Lp_2162, SP_3050, His_6_ tag | This work |
| pJRB03Q-LSH | pSIP401 based plasmid containing Ls-araA, sppQ, Lp_2940, SP_3050, His_6_ tag | This work |
| pJRB04Q-LSH | pSIP401 based plasmid containing Ls-araA, sppQ, His_6_ tag | This work |
| pJRB05Q-LSH | pSIP401 based plasmid containing Ls-araA, sppQ, Lp_1261, SP_3050, His_6_ tag | This work |
| pJRB06Q-LSH | pSIP401 based plasmid containing Ls-araA, sppQ, Lp_1452, SP_3050, His_6_ tag | This work |
| pJRB08Q-LSH | pSIP401 based plasmid containing Ls-araA, sppQ, Lp_3014, SP_3050, His_6_ tag | This work |
| pJRB09Q-LSH | pSIP401 based plasmid containing Ls-araA, SP_3050, His_6_ tag | This work |
| pJRB14Q-LSH | pSIP411 based plasmid containing Ls-araA, sppQ, His_6_ tag | This work |

**Supplementary Table 3: Strains used in this study**

| **Strain** | **Description** |
| --- | --- |
| *E. coli* NEB 5α | NEB (Beverly, MA) |
| *L. plantarum* WCFS1 | NIZO Food Research (Kernhemseweg, Netherlands) |
| A1 | *L. plantarum* containing plasmid pJRB01Q-LSH |
| A2 | *L. plantarum* containing plasmid pJRB02Q-LSH |
| A3 | *L. plantarum* containing plasmid pJRB03Q-LSH |
| A4 | *L. plantarum* containing plasmid pJRB05Q-LSH |
| A5 | *L. plantarum* containing plasmid pJRB06Q-LSH |
| A6 | *L. plantarum* containing plasmid pJRB08Q-LSH |
| SEC | *L. plantarum* containing plasmid pJRB09Q-LSH |
| IC1 | *L. plantarum* containing plasmid pJRB04Q-LSH |
| IC2 | *L. plantarum* containing plasmid pJRB14Q-LSH |
| IC2 + PBS | Unmodified strain IC2 |
| IC2 + SDS | Modified strain IC2 permeabilized with 0.01% SDS |

**Supplementary Table 4: Native *L. plantarum* anchor proteins used for LsLAI surface display.**

| **Strain** | **Anchor protein** | **Orientation** | **Type** | **Ref.** |
| --- | --- | --- | --- | --- |
| **A1** | Lp_2578 | C-terminal | LPxTG | ^1,2^ |
| **A2** | Lp_2162 | C-terminal | LysM | ^3^ |
| **A3** | Lp_2940 | C-terminal | LPxTG | ^4^ |
| **A4** | Lp_1261 | N-terminal | Lipobox | ^2,5^ |
| **A5** | Lp_1452 | N-terminal | Lipobox | ^6^ |
| **A6** | Lp_3014 | N-terminal | LysM | ^6,7^ |

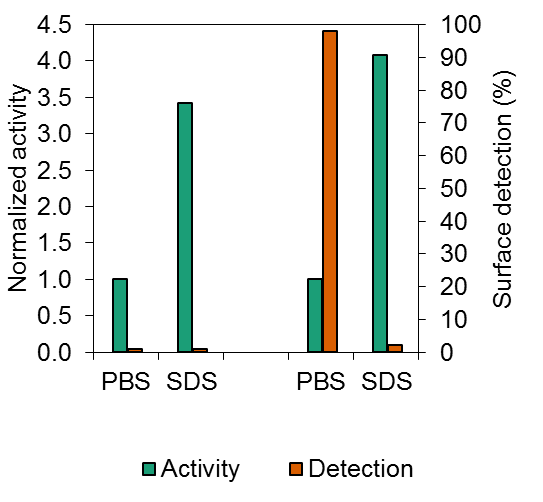

Intracellular

Surface Display

**Supplementary Figure 1: Surface treatment of *L. plantarum* surface displayed LsLAI.** a) Comparison of the activity (green) and surface detection percentage (orange) of *L. plantarum* expressing LsLAI containing a His_6_-tag either intracellularly (left) or surface displayed with anchor protein A6 (right). Activity of cells treated with 0.05 % SDS normalized to untreated cells (PBS). b) Western blot analysis of insoluble (lanes 2 - 4) or soluble protein fraction (lanes 5 - 6) of *L. plantarum* wild-type “WT” (lanes 2, 5) or expressing LsLAI intracellularly “IC1” (lanes 3, 6) or expressing A6-LsLAI surface displayed “SD” (lanes 4, 7). Expected molecular weight (MW) of LsLAI and A6-LsLAI is 54 kDa and 76.5 kDa, respectively.

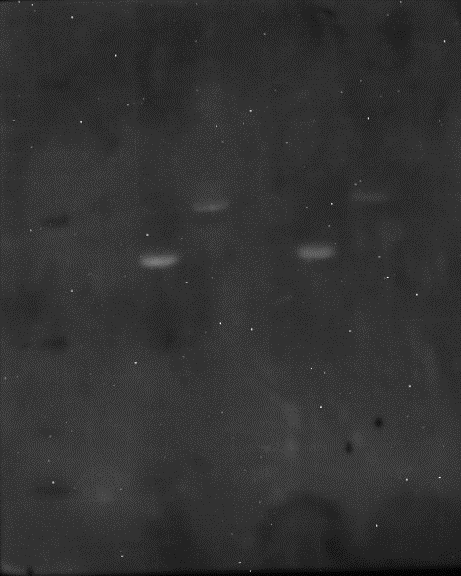

225

115

80

65

50

35

30

25

15

10

1

2

3

4

5

6

7

Lane

MW (kDa)

LsLAI

A6-LsLAI

Ladder

Insoluble

Soluble

WT

WT

IC1

SD

IC1

SD

**a**

**b**

Normalized activity

Surface detection (%)

|  | Activity | Detection |
| --- | --- | --- |
| PBS | 1 | 1 |
| SDS | 3.42857 | 1 |
| PBS | 1 | 98 |
| SDS | 4.08333 | 2 |

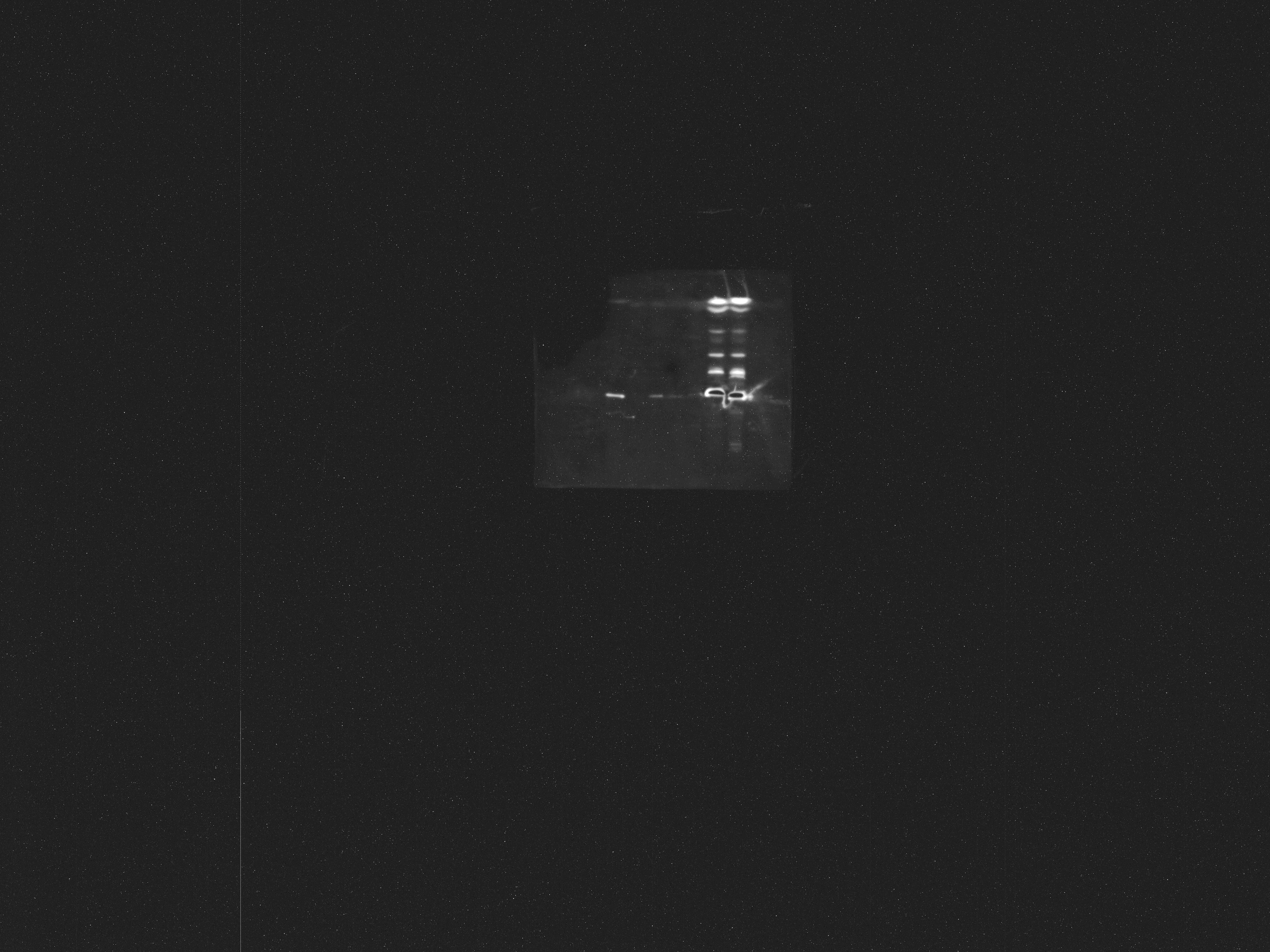

225

80

65

50

30

10

MW (kDa)

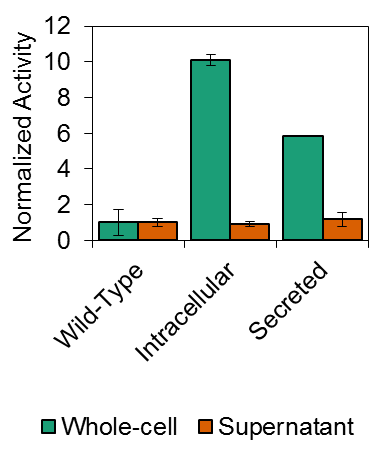

Lane

1

2

3

4

5

6

7

8

9

10

Ladder

Supernatant

Soluble

WT

IC1

Sec

Insoluble

WT

IC1

Sec

WT

IC1

Sec

**a**

**b**

**Supplementary Figure 2: *L. plantarum* secreted LsLAI is inactive.** a) Comparison of the whole-cell (green) and supernatant (orange) of *L. plantarum* wild-type (left) or expressing LsLAI containing a His_6_-tag either intracellularly or as secreted/unanchored protein. b) Western blot analysis of culture supernatant (lanes 2 - 4) or soluble (lanes 5 - 7) and insoluble (lanes 8 - 10) protein fractions of *L. plantarum* cells. Shown are wild-type “WT” control cells (lanes 2, 5, 8), cells expressing LsLAI intracellularly “I” (lanes 3, 6, 9), and cells secreting LsLAI “Sec” (lanes 4, 7, 10). Supernatant was concentrated 20 × before analysis. Expected molecular weight (MW) of LsLAI and secreted LsLAI is 54 kDa.

LAI

Normalized Activity

|  | Whole-cell | Supernatant |
| --- | --- | --- |
| Wild-Type | 1 | 1 |
| Intracellular | 10.1014 | 0.909743 |
| Secreted | 5.82374 | 1.17334 |

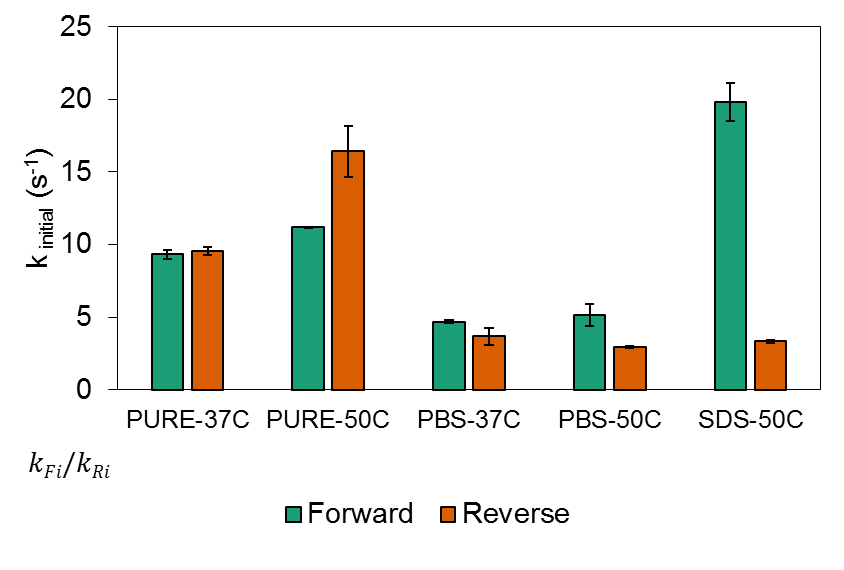

a

b

a,c

b,d,e

f

c,f

e

d

1.0

0.7

1.3

1.8

5.9

**Supplementary Figure 3: Initial turnover rates.** Initial turnover rates of purified free-enzyme, (PURE), *L. plantarum*-encapsulated (PBS), and permeabilized *L. plantarum* encapsulated (SDS) LsLAI in forward (galactose as substrate) (green) and reverse (tagatose as substrate) direction (orange) in the presence of 400 mM substrate at 37 or 50 °C. Ratio of forward to reverse reaction rate is denoted by k_Fi_/k_Ri_. The data are means from three biological replicates. (Significance between samples tested via ANOVA analysis using SigmaPlot 13.0. a,c,d,e = p < 0.001, b = p < 0.05)

k initial (s-1)

|  | Forward | Reverse |
| --- | --- | --- |
| PURE-37C | 9.32678 | 9.537 |
| PURE-50C | 11.1853 | 16.4208 |
| PBS-37C | 4.69912 | 3.68843 |
| PBS-50C | 5.14471 | 2.92297 |
| SDS-50C | 19.8165 | 3.33134 |

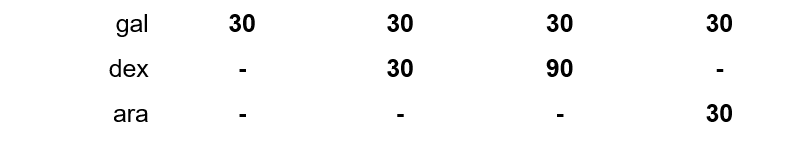

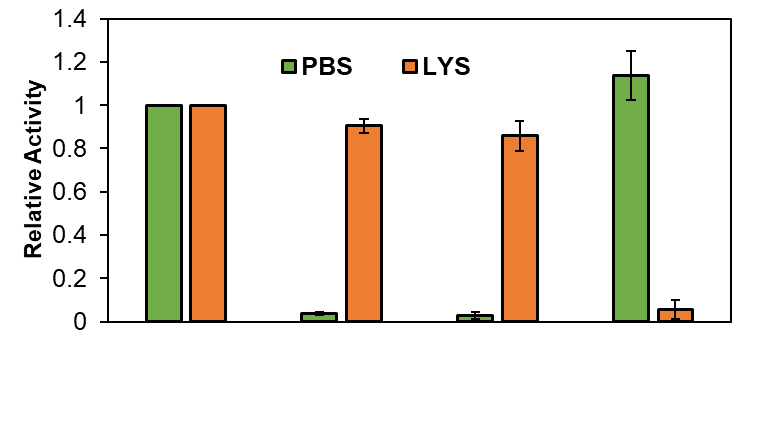

**Supplementary Figure 4: Selective nature of cellular encapsulating LsLAI**. Tagatose production of unmodified IC2 cells expressing LsLAI (PBS, green) or cell lysate (LYS, orange) in the presence of different combinations of galactose (gal), dextrose (dex), and/or arabinose (ara) after 20 min incubation at 37 °C. Activity normalized to 30 mM galactose condition for unmodified whole-cells (PBS) and cell lysate (LYS) independently. The data are means from three biological replicates.

**Concentration (mM)**

Relative Activity

|  | PBS | LYS |
| --- | --- | --- |
| 0:1 | 1 | 1 |
| 1:1 | 0.0367332 | 0.903969 |
| 3:1 | 0.0273226 | 0.858301 |
| Arabinose | 1.13684 | 0.0547546 |

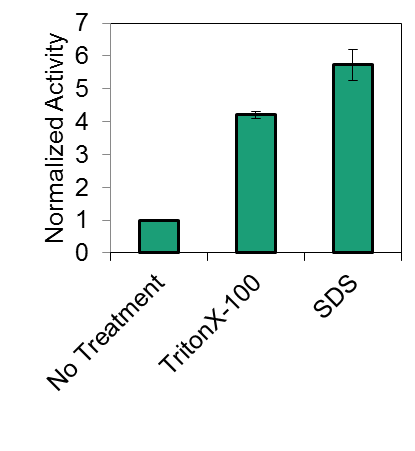

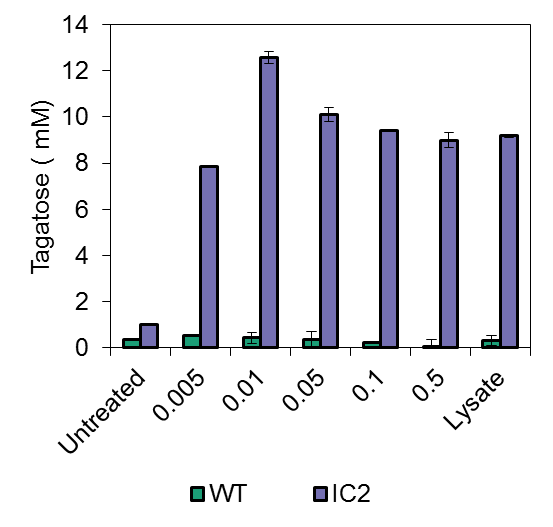

SDS (%)

1%

0.01%

**CMC at 37 °C**

**Concentration**

0.05%

0.25%

+

-

**a**

**b**

**Supplementary Figure 5: SDS permeabilization of encapsulated LsLAI overcomes kinetic penalty.** a, Comparison of activity of encapsulated LsLAI having undergone permeabilization by 1 % TritonX-100 (middle) or 0.01 % SDS (right) normalized to untreated cells “No Treatment” (left). b, Optimization of SDS permeabilization of *L. plantarum* wild type “WT” (green) or expressing LsLAI intracellularly “IC2” (purple) that produced the greatest amount of tagatose at 37 °C in 2 h as compared to untreated cells or crude lysate. The data are means from three biological replicates.

|  | Normalized Activity |
| --- | --- |
| No Treatment | 1 |
| TritonX-100 | 4.21227 |
| SDS | 5.73431 |

Tagatose ( mM)

|  | WT | IC2 |
| --- | --- | --- |
| Untreated | 0.353135 | 1.01171 |
| 0.005 | 0.534545 | 7.8522 |
| 0.01 | 0.435254 | 12.5837 |
| 0.05 | 0.387588 | 10.1121 |
| 0.1 | 0.229704 | 9.40433 |
| 0.5 | 0.0628838 | 8.99732 |
| Lysate | 0.322651 | 9.17677 |

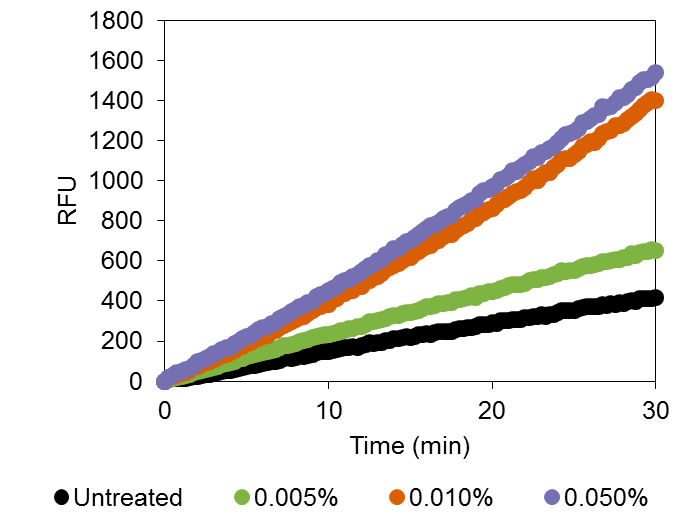

SDS

14.3

22.4

44.3

49.2

RFU/min

**Supplementary Figure 6: SDS permeabilization increases transport.** Continuously monitoring the production of fluorescent signal from reporter cFDA after activation upon transport using *L. plantarum* wild-type untreated (black) or after SDS permeabilization with 0.005 % (green), 0.01 % (orange), or 0.05 % (purple) SDS. Relative fluorescence units (RFU) generated per minute is a proxy for transport kinetics.

RFU

| Time (min) | Untreated | 0.005% | 0.010% | 0.050% |
| --- | --- | --- | --- | --- |
| 0 | 0 | 0 | 0 | 0 |
| 0.25 | 5.88 | 9.848 | 16.882 | 18.767 |
| 0.5 | 14.151 | 10.624 | 21.436 | 28.512 |
| 0.75 | 12.839 | 17.547 | 29.514 | 42.274 |
| 1 | 10.9 | 23.674 | 39.757 | 48.853 |
| 1.25 | 13.892 | 26.74 | 43.639 | 57.942 |
| 1.5 | 18.801 | 31.419 | 53.306 | 64.592 |
| 1.75 | 22.243 | 42.484 | 66.826 | 79.63 |
| 2 | 22.783 | 49.392 | 74.661 | 95.654 |
| 2.25 | 28.245 | 50.97 | 84.606 | 101.108 |
| 2.5 | 34.043 | 59.509 | 90.442 | 111.704 |
| 2.75 | 32.596 | 63.797 | 97.589 | 121.526 |
| 3 | 39.889 | 69.161 | 109.747 | 136.542 |
| 3.25 | 44.348 | 74.831 | 117.024 | 143.869 |
| 3.5 | 48.108 | 82.105 | 128.49 | 157.666 |
| 3.75 | 51.301 | 87.368 | 134.907 | 166.85 |
| 4 | 51.747 | 92.827 | 144.392 | 169.618 |
| 4.25 | 58.401 | 98.267 | 153.623 | 187.488 |
| 4.5 | 64.692 | 113.665 | 165.45 | 198.87 |
| 4.75 | 67.42 | 113.151 | 171.508 | 210.584 |
| 5 | 72.102 | 121.266 | 189.914 | 223.663 |
| 5.25 | 76.369 | 128.204 | 192.509 | 228.767 |
| 5.5 | 79.489 | 131.175 | 207.404 | 242.816 |
| 5.75 | 82.208 | 137.049 | 214.651 | 257.539 |
| 6 | 89.196 | 144.444 | 224.541 | 266.634 |
| 6.25 | 92.982 | 145.748 | 226.322 | 274.307 |
| 6.5 | 96.82 | 154.893 | 245.147 | 286.873 |
| 6.75 | 98.641 | 159.976 | 251.75 | 294.346 |
| 7 | 103.003 | 167.337 | 265.823 | 310.96 |
| 7.25 | 110.346 | 175.27 | 275.93 | 323.012 |
| 7.5 | 117.884 | 176.277 | 286.192 | 334.178 |
| 7.75 | 114.386 | 183.039 | 295.028 | 347.834 |
| 8 | 118.22 | 188.148 | 306.55 | 354.899 |
| 8.25 | 126.275 | 197.417 | 319.508 | 370.603 |
| 8.5 | 125.042 | 202.382 | 323.562 | 379.34 |
| 8.75 | 128.907 | 206.05 | 337.395 | 390.703 |
| 9 | 135.067 | 217.308 | 350.041 | 399.19 |
| 9.25 | 142.677 | 220.081 | 365.697 | 420.026 |
| 9.5 | 146.709 | 227.446 | 369.833 | 432.78 |
| 9.75 | 147.84 | 230.985 | 382.826 | 436.714 |
| 10 | 149.197 | 233.338 | 383.531 | 449.86 |
| 10.25 | 152.146 | 239.212 | 406.42 | 462.131 |
| 10.5 | 156.871 | 242.782 | 410.422 | 474.481 |
| 10.75 | 162.716 | 254.262 | 429.912 | 489.855 |
| 11 | 165.566 | 254.102 | 436.712 | 500.795 |
| 11.25 | 169.591 | 262.296 | 451.426 | 508.057 |
| 11.5 | 173.037 | 264.799 | 453.014 | 523.741 |
| 11.75 | 167.913 | 271.438 | 470.544 | 532.65 |
| 12 | 178.226 | 272.003 | 474.543 | 549.014 |
| 12.25 | 183.892 | 282.874 | 495.923 | 559.555 |
| 12.5 | 191.694 | 296.314 | 503.759 | 575.649 |
| 12.75 | 188.341 | 296.058 | 518.97 | 582.142 |
| 13 | 192.977 | 300.244 | 523.459 | 599.479 |
| 13.25 | 193.299 | 306.416 | 539.292 | 608.212 |
| 13.5 | 203.643 | 314.239 | 557.778 | 631.091 |
| 13.75 | 201.639 | 320.2 | 564.554 | 634.339 |
| 14 | 210.229 | 326.197 | 576.785 | 659.675 |
| 14.25 | 211.685 | 325.764 | 587.218 | 661.57 |
| 14.5 | 216.301 | 335.849 | 600.868 | 683.424 |
| 14.75 | 220.982 | 342.786 | 617.933 | 694.957 |
| 15 | 219.226 | 342.396 | 617.687 | 700.935 |
| 15.25 | 221.906 | 347.614 | 636.41 | 717.835 |
| 15.5 | 234.093 | 351.658 | 647.03 | 731.905 |
| 15.75 | 233.369 | 362.322 | 663.229 | 745.294 |
| 16 | 237.41 | 370.069 | 671.127 | 762.624 |
| 16.25 | 234.902 | 373.574 | 676.818 | 774.11 |
| 16.5 | 240.011 | 375.078 | 694.479 | 779.81 |
| 16.75 | 245.102 | 385.955 | 702.46 | 791.472 |
| 17 | 245.382 | 385.168 | 721.032 | 813.345 |
| 17.25 | 247.486 | 392.572 | 734.867 | 821.633 |
| 17.5 | 249.694 | 394.513 | 731.888 | 828.558 |
| 17.75 | 258.767 | 406.146 | 752.689 | 849.043 |
| 18 | 263.847 | 406.46 | 764.497 | 866.518 |
| 18.25 | 264.011 | 410.318 | 775.024 | 874.736 |
| 18.5 | 265.488 | 416.977 | 786.948 | 886.021 |
| 18.75 | 269.507 | 426.875 | 804.782 | 900.544 |
| 19 | 270.755 | 418.622 | 809.929 | 905.851 |
| 19.25 | 283.982 | 434.934 | 831.451 | 934.123 |
| 19.5 | 279.925 | 437.222 | 842.051 | 949.93 |
| 19.75 | 284.608 | 446.03 | 854.401 | 954.047 |
| 20 | 291.508 | 449.291 | 861.371 | 970.154 |
| 20.25 | 285.808 | 454.324 | 879.309 | 982.52 |
| 20.5 | 300.224 | 461.305 | 903.616 | 1004.85 |
| 20.75 | 301.834 | 468.762 | 908.045 | 1012.65 |
| 21 | 303.027 | 475.098 | 921.948 | 1025.35 |
| 21.25 | 307.914 | 481.716 | 937.93 | 1053.38 |
| 21.5 | 307.545 | 480.15 | 948.416 | 1047.18 |
| 21.75 | 309.205 | 484.333 | 953.687 | 1064.82 |
| 22 | 316.932 | 488.222 | 967.771 | 1081.53 |
| 22.25 | 318.703 | 503.001 | 992.469 | 1095.61 |
| 22.5 | 322.759 | 508.153 | 1010.06 | 1119.37 |
| 22.75 | 320.605 | 507.153 | 1002.94 | 1115.93 |
| 23 | 331.516 | 516.534 | 1024.19 | 1141.54 |
| 23.25 | 323.388 | 516.759 | 1040.74 | 1145.08 |
| 23.5 | 332.78 | 522.008 | 1039.98 | 1162.14 |
| 23.75 | 333.95 | 529.96 | 1066.31 | 1173.33 |
| 24 | 341.012 | 536.183 | 1078.72 | 1190.77 |
| 24.25 | 351.538 | 553.871 | 1108.04 | 1212.71 |
| 24.5 | 350.804 | 549.018 | 1111.49 | 1228.3 |
| 24.75 | 354.275 | 553.505 | 1110.68 | 1236.98 |
| 25 | 353.028 | 554.057 | 1131.52 | 1244.05 |
| 25.25 | 362.898 | 563.005 | 1148.28 | 1263.12 |
| 25.5 | 367.653 | 569.812 | 1170.81 | 1289.35 |
| 25.75 | 371.66 | 577.218 | 1179.03 | 1298 |
| 26 | 370.232 | 578.745 | 1195.46 | 1311.05 |
| 26.25 | 372.55 | 580.718 | 1191.88 | 1324.83 |
| 26.5 | 374.146 | 584.744 | 1209.76 | 1329.36 |
| 26.75 | 383.172 | 598.648 | 1236.84 | 1368.98 |
| 27 | 378.602 | 596.795 | 1245.47 | 1363.06 |
| 27.25 | 387.075 | 603.285 | 1249.44 | 1372.26 |
| 27.5 | 384.187 | 606.896 | 1274.24 | 1390.67 |
| 27.75 | 390.57 | 606.877 | 1273.84 | 1414.64 |
| 28 | 389.146 | 616.435 | 1287.47 | 1413 |
| 28.25 | 396.745 | 615.358 | 1306.51 | 1432.9 |
| 28.5 | 401.063 | 636.37 | 1320.45 | 1455.26 |
| 28.75 | 399.09 | 629.077 | 1334.69 | 1464.63 |
| 29 | 411.004 | 640.301 | 1353.21 | 1490.79 |
| 29.25 | 411.001 | 645.891 | 1374.26 | 1503.24 |
| 29.5 | 412.933 | 645.168 | 1388.83 | 1502.96 |
| 29.75 | 411.958 | 658.961 | 1406.59 | 1516.67 |
| 30 | 414.859 | 651.131 | 1402.64 | 1537.99 |

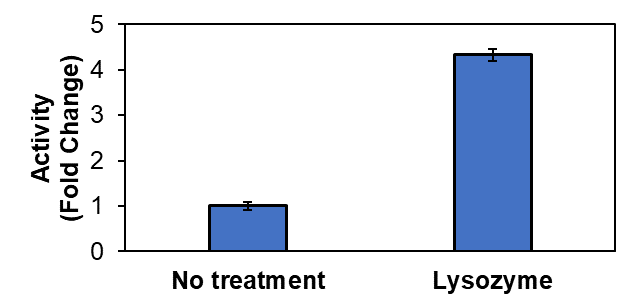

**Supplementary Figure 7: Lysozyme treated cells have enhanced tagatose production.** Comparing tagatose production of encapsulated LsLAI (IC2) having undergone treatment with 0.01 μM lysozyme (right) normalized to untreated cells “No Treatment” (left) in the presence of 200 mM galactose after 2 h incubation at 37 °C. The data are means from three biological replicates.

|  | Activity (Fold Change) |
| --- | --- |
| No treatment | 1 |
| Lysozyme | 4.32773 |

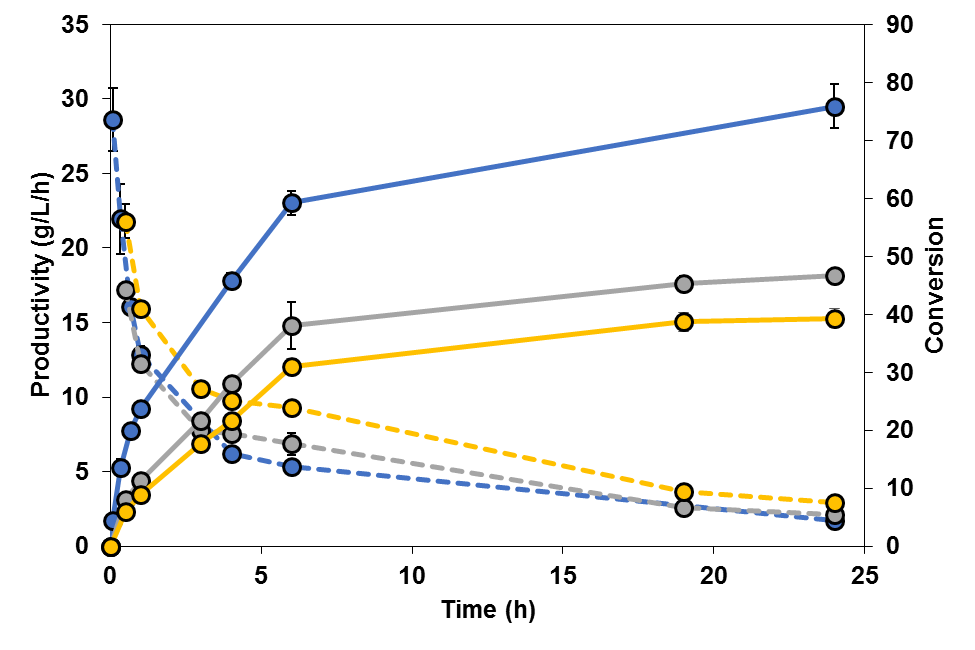

**Supplementary Figure 8: Comparing conversion and productivity of SDS treated encapsulated LsLAI at different initial galactose concentrations.** Measuring the conversion and average productivity of tagatose production using permeabilized encapsulated LsLAI incubated at 50 °C “SDS-50C” at 300 mM galactose (blue), 600 mM galactose (silver), or 1 M galactose (yellow). Average productivity calculated at each sample timepoint. Inset shows initial reaction rates. Data for 300 mM galactose taken from Figure 4 of this work. The data are means from three biological replicates.

Conversion

| Time (h) | Productivity (g/L/h) |
| --- | --- |
| 0.0833333 | 28.6122 |
| 0.333333 | 21.9457 |
| 0.666667 | 16.0901 |
| 1 | 12.8637 |
| 4 | 6.18444 |
| 6 | 5.32857 |
| 24 | 1.7079 |

| Time (h) |  |  |
| --- | --- | --- |
| 0.5 | 17.2193 | 21.7985 |
| 1 | 12.2589 | 15.9692 |
| 3 | 7.77703 | 10.6023 |
| 4 | 7.56548 | 9.75654 |
| 6 | 6.8503 | 9.3076 |
| 19 | 2.57519 | 3.66367 |
| 24 | 2.10453 | 2.9481 |

|  | 300mM-Conv |
| --- | --- |
| 0 | 0 |
| 0.0833333 | 4.41546 |
| 0.333333 | 13.5467 |
| 0.666667 | 19.8643 |
| 1 | 23.8217 |
| 4 | 45.8107 |
| 6 | 59.2063 |
| 24 | 75.9066 |

|  | 600mM-Conv | 1M-Conv |
| --- | --- | --- |
| 0 | 0 | 0 |
| 0.5 | 7.97191 | 6.05515 |
| 1 | 11.3509 | 8.87177 |
| 3 | 21.6029 | 17.6706 |
| 4 | 28.0203 | 21.6812 |
| 6 | 38.0572 | 31.0253 |
| 19 | 45.3043 | 38.672 |
| 24 | 46.7672 | 39.308 |

**Supplementary Figure 9: TEM analysis of SDS permeabilized *L. plantarum*.** Transmission electron microscopy of *L. plantarum* cells to study the effects of 0.01 %SDS permeabilization on cellular structure. a, Wild-type *L. plantarum* in PBS. b, Strain IC2 intracellularly expressing LsLAI in PBS. c, Strain IC2 intracellularly expressing LsLAI treated with 0.01% SDS. HV = 80.0 kV. Direct Mag: 49000×.

**a**

**c**

**b**

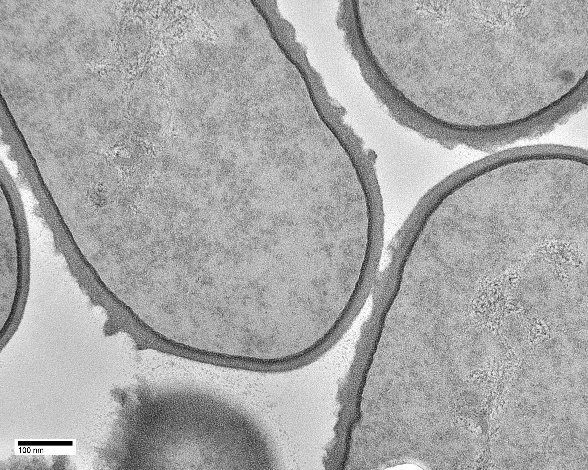

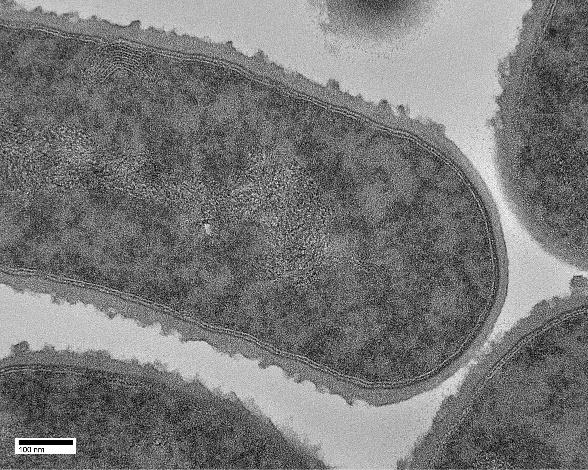

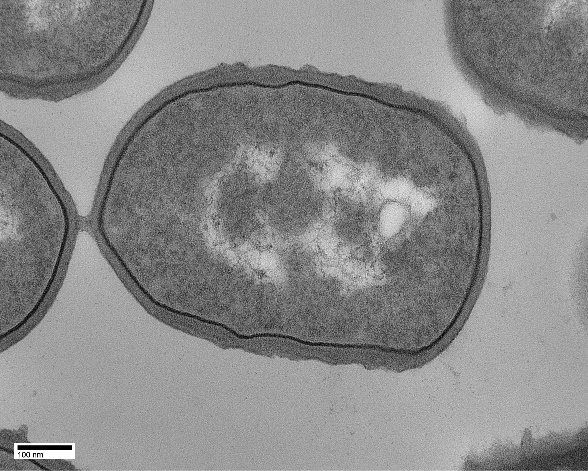

**Supplementary Figure 11: HPLC chromatograms of tagatose production samples using strain IC2 + SDS at 50 °C.** ELSD chromatogram signals at 0, 6, 24, and 48 h timepoints. One representative figure of triplicate samples.

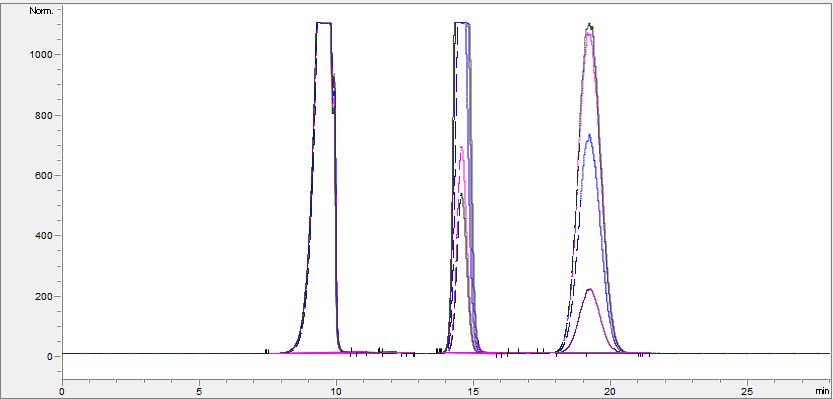

**galactose**

**tagatose**

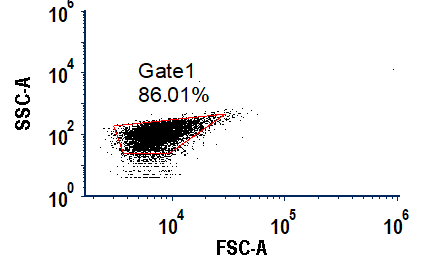

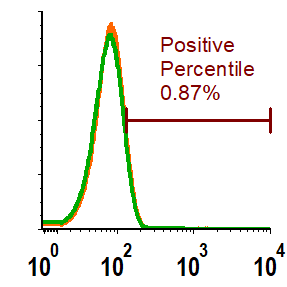

**Supplementary Figure 10: Flow cytometry analysis gating.** a) Population gating of the negative control (wild-type). Axes are bio-exponential. b) Marker of the negative controls wild-type and intracellularly expressed LsLAI with His_6_-tag. Histogram counts normalized. Plotted on bio-exponential x-axis. Positive percentile marker is shown.

**a**

**b**
